# Evolutionary Dynamics of Lineage-Specific Class-A GPCR Subsets Reveal Widespread Chemosensory Roles and Adaptations in Lophotrochozoa

**DOI:** 10.1101/2024.07.14.603414

**Authors:** Rohan Nath, Biswajit Panda, Rakesh Siuli, Arunkumar Krishnan

## Abstract

Detecting external chemosensory cues via Class-A G protein-coupled receptors (GPCRs) is essential for behavioral and biological functions, influencing animal evolution and ecological adaptations. While well-studied in vertebrates and echinoderms, their role in major clades like Lophotrochozoa is less understood despite their remarkable ecological adaptations. Utilizing 238 lophotrochozoan genomes across eight phyla, we conducted a large-scale comparative genomics analysis to identify lineage-specifically expanded subsets (LSEs) of Class-A GPCRs adapted for chemoreception. Using phylogeny and orthology-based clustering, we differentiated these expansions from conserved orthogroups of endogenous ligand-binding GPCRs. LSEs correlated with adaptations to diverse habitats, with whole-genome duplications having limited impact. Across phyla, species in coastal, freshwater, and terrestrial habitats exhibited large and diverse LSEs, while those adapted to extreme deep-sea environments, parasitic lifestyles, or alternative chemosensory mechanisms showed consistent reductions. Sequence heterogeneity, positive selection, and ligand-binding pocket flexibility in these LSEs further underscored adaptations to environmental signals. These findings provide foundational insights into Class-A GPCR-mediated chemoreception across Lophotrochozoa.

**Teaser:** Unveiling correlations between lophotrochozoans habitat adaptations and lineage-specific changes in Class-A GPCR repertoire.

## INTRODUCTION

Sensing the dynamic external milieu or chemosensory cues via cell-surface proteins is essential for a multitude of behavioral and biological functions and plays a fundamental role in the evolution and ecological adaptations of metazoans (*1–3*). Multiple chemoreceptor families (CRs) have independently evolved in metazoans to coordinate external chemosensory cues, eliciting various physiological responses ranging from larval settlement to activities like finding mates, reproduction, navigation, predator avoidance, and a range of other neuronal functions across diverse phyla (*2, 4–8*). Extensively studied CRs are categorized into distinct families (*2*), and of these, the 7TM Class-A G protein-coupled receptor (GPCR) family, comprising the vertebrate olfactory receptors (ORs) (*9*), is notable for exhibiting extensive lineage-specific expansions (LSEs) with significant interspecific variation in gene counts, driven by gene duplications, losses, and pseudogenization (*2, 3, 10–13*). These expansions mostly correlate with differences in habitat, diet, and environment, emphasizing ecological influences on their evolution (*13*).

While distinct evolutionary patterns among CR families show an increased prevalence of ionotropic receptors in protostomes and more metabotropic GPCR-mediated chemoreception in vertebrates (*14–18*), previous investigations have, however, identified LSEs of chemosensory Class-A GPCRs beyond vertebrates in both deuterostome and protostome lineages. These are exemplified by LSEs of Class-A GPCRs in (i) *Strongylocentrotus purpuratus* and *Acanthaster planci*, which are particularly enriched in external and sensory tissues (*19–21*); (ii) in *Caenorhabditis elegans* (*22, 23*); (iii) in gastropod mollusks like *Aplysia californica*, with differential expression in rhinophores and oral tentacles (*24, 25*).

These findings collectively underscore the following: (i) the role of Class-A GPCRs in chemoreception is more widespread across diverse invertebrate groups than previously recognized (*19–21, 23–25*); (ii) Class-A GPCRs adapted for chemosensory roles specifically exhibit diverse LSEs (*19–21, 24*), whereas the endogenous ligand-binding subfamilies form respective orthogroups comprising conserved one-to-one orthologs from various lineages. These insights raise the question of what proportion of Class-A GPCRs exhibits diverse LSEs in various previously overlooked invertebrate phyla and how these clusters correlate with diverse ecological factors.

To investigate these inquiries and comprehend the evolutionary dynamics of Class-A GPCR LSEs within a vastly overlooked invertebrate clade demonstrating remarkable adaptability across aquatic and terrestrial environments, we here consider Lophotrochozoa as an exemplary case study. Previous investigations have largely been confined to *Aplysia californica* or limited to no more than three distinct taxa within the subclass caenogastropods (*24–26*). With the availability of over 230 genomes spanning various phyla, the clade Lophotrochozoa, presents a prime opportunity to study the evolution and diversifications of chemosensory GPCRs and how adaptations to different environments have shaped the repertoire of these receptors. Using a multi-pronged approach, we here, for the first time, classified the LSEs of Class-A GPCRs in 238 genomes and provided unprecedented details by delineating the highly variable trends in the interspecific variations of these receptors across Lophotrochozoa.

## RESULTS

### Phylogeny and Orthology-based Clustering Approaches Identify and Distinguish Class-A GPCR LSEs

Utilizing a comprehensive comparative genomics approach, we initially curated the Class-A GPCR repertoire from 238 genomes spanning 8 distinct phyla of Lophotrochozoa. Following this, we employed an integration of phylogenetic analysis and OrthoFinder-based methods to identify and classify lineage-specifically expanded subsets distinct from conserved orthogroups of prospective endogenous ligand-binding counterparts. A hierarchical and recursive approach to phylogenetic tree reconstruction across various taxonomic levels and combinations was utilized to precisely delineate the taxa-specific paralogous clusters.

Additionally, OrthoFinder was utilized to segregate sequences that did not converge into orthogroups, and LSEs identified through both methodologies were cross-validated. Specifically, LSEs identified through phylogenetic analysis underwent scrutiny with OrthoFinder to confirm the absence of orthologous relationships and vice versa. The validated LSEs were then subjected to further analysis, including assessment of tandem clustering patterns in the genome, and all validated sequences forming LSEs were categorized into full-length, truncated, and pseudogenes. The overall pipeline is presented as a flowchart (Fig. 1A), with an elaborate description summarized in the methods. Thus, the Class-A GPCR and corresponding LSE counts of GPCRs were derived from each examined taxon, and Fig. 1B presents their total count, percentage proportion within the Class-A GPCR complement, and counts of full-length, truncated, and pseudogenes for a representative list of analyzed taxa. In total, we classified 108,822 Class-A GPCRs across 238 genomes: of these, 46,798 were categorized as LSEs, while 62,024 belonged to conserved orthogroups. The distribution of LSE proportions reveals that 52 genomes have over 50% of their Class-A GPCRs as LSEs. Notably, 8 genomes fall within the 60-70% range, 9 genomes within the 70-80% range, and 5 genomes encode extreme expansions as over 80% of the Class-A GPCR subset are LSEs. Among the genomes with less than 50% as LSEs, 69 genomes exhibit over 30%, and 29 show over 40% (table S1).

**Fig. 1.**
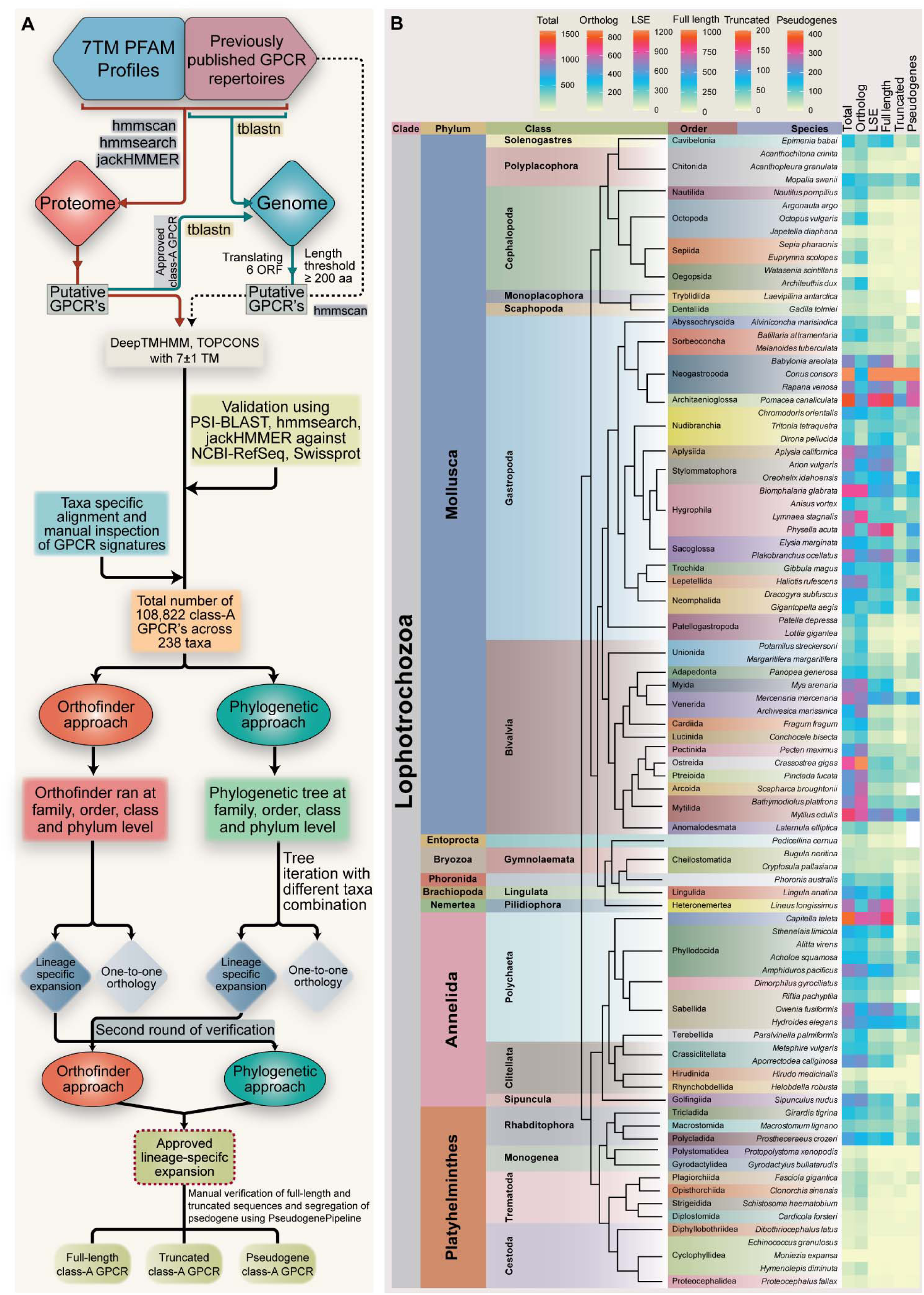
Overview of the LSE identification protocol and summary of Class-A GPCR LSEs in Lophotrochozoa. **(A)** Schematic illustration of the methodological workflow used for the systematic identification of Class-A GPCR LSEs across each analyzed species. **(B)** Summary of the representative list of analyzed taxa, their taxonomic classifications, and gene counts. The heatmap on the right depicts expansions and contractions of total Class-A GPCR counts, including one-to-one orthologs and LSE subsets. Identified LSEs are categorized into full-length, truncated, and pseudogene counts, with the color scale for each heatmap column displayed at the top.

In summary, significant expansions are observed in the majority of the genomes analyzed in Gastropoda (Class-A range: 226-1560 genes, mean: 689; LSEs: 25-1226, mean: 370; 46 species analyzed), followed by Bivalvia (Class-A range: 264-1266 genes, mean: 674; LSEs: 10-644, mean: 226; 60 species analyzed), and Annelida (Class-A range: 8-1481 genes, mean: 505; LSEs: 0-889, mean: 245; 32 species analyzed) with specific exceptions that correlate with the ecological and lifestyle adaptations (table S1). Other molluscan classes (Class-A range: 140-527 genes, mean: 303; LSEs: 31-350, mean: 142; 9 species analyzed) and other lophotrochozoan phyla (Class-A range: 123-2352 genes, mean: 515; LSEs: 24-1904, mean: 352; 13 species analyzed) also show notable expansions. In contrast, Cephalopoda (Class-A range: 46-411 genes, mean: 183; LSEs: 3-146, mean: 29; 19 species analyzed) and Platyhelminthes (Class-A range: 2-704 genes, mean: 128; LSEs: 0-370, mean: 32; 59 species analyzed) exhibit considerable reductions (table S1). We found a significant difference among the various groups using the Kruskal-Wallis rank sum test (Class-A: P = 2.2x10^-16^; LSEs: P = 2.2x10^-16^), followed by conducting Dunn’s post-hoc test with Bonferroni correction for pairwise comparisons (Fig. 2A and B). Overall, the results of our study reveal unique expansions and contractions of the candidate chemosensory GPCRs that significantly align with factors such as lifestyle, ecological niche, and specific adaptations. Subsequent sections present a detailed elucidation of these trends for each phylum within Lophotrochozoa.

**Fig. 2.**
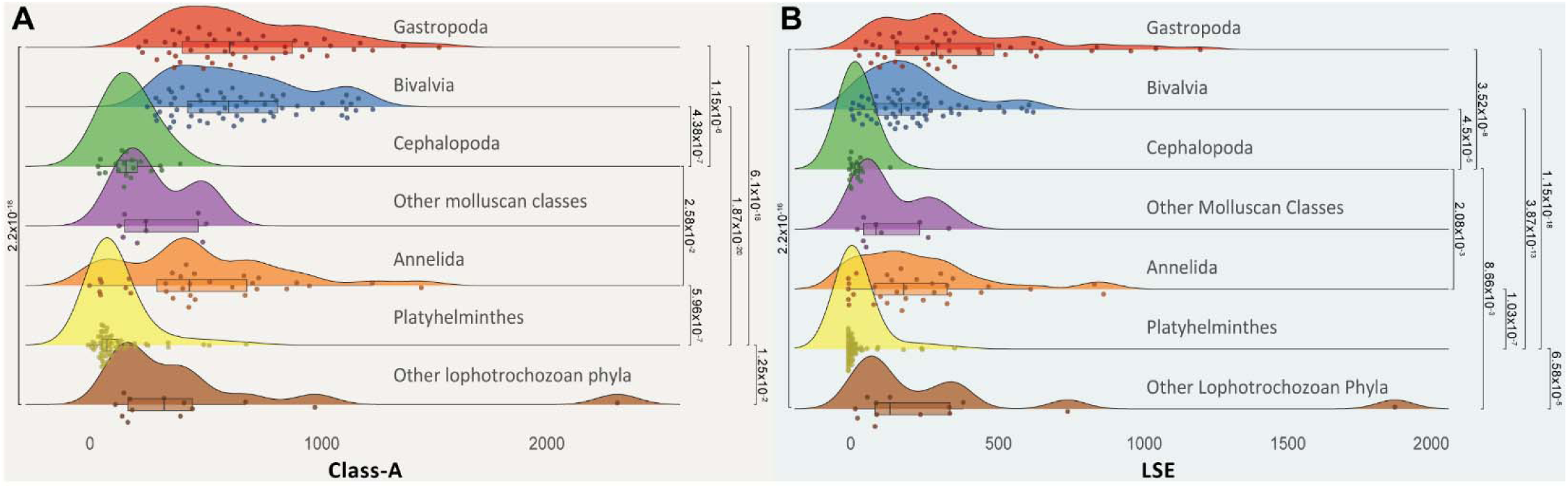
Significant variations in Class-A GPCR repertoire and LSE counts across groups. Raincloud plots illustrate statistically significant pair-wise comparisons among analyzed groups for overall Class-A GPCR counts **(A)** and LSE subsets **(B)**. Statistical significance was assessed using the Kruskal-Wallis rank sum test, followed by Dunn’s post-hoc analysis with Bonferroni correction. Only significant differences (p < 0.05) are reported. Other molluscan classes include Solenogastres, Polyplacophora, Monoplacophora, and Scaphopoda, while additional lophotrochozoan phyla include Entoprocta, Bryozoa, Phoronida, Brachiopoda, and Nemertea.

### Pan-Genomic Insights into Candidate Chemosensory GPCRs Across Mollusca

As the second most species-rich phylum in the animal kingdom, mollusks have flourished in both aquatic and terrestrial environments, showcasing notable adaptations mirrored in gene-family innovations (*27, 28*). Our comprehensive pan-genomic analysis, involving 134 molluscan species across seven classes, revealed intriguing trends.

#### Overall patterns in gastropods and bivalves

Examining the genomes of 46 gastropod species spanning 13 orders reveals that, on average, close to half of the Class-A GPCR repertoire are LSEs (Fig. 3A-C). Consistent with earlier observations (*25*), certain marine gastropods exhibit notably high proportions; for instance, the *Conus* and *Pomacea* genera encode close to a thousand genes as LSEs, representing some of the most extensive expansions with a significant difference compared to other lophotrochozoans (*Conus* LSEs: 865-1226; *Pomacea* LSEs: 984; P = 7.03x10^-4^, Wilcoxon rank sum with continuity correction) (Fig. 3D). Analysis of 60 bivalve species across 12 orders, indicates a significant prevalence of LSEs, accounting for around 30% of Class-A GPCRs on average, with most bivalve orders displaying notable expansions (Fig. 4A-C). However, gastropods and bivalves adapted to extreme environments show reduced proportions of Class-A GPCRs and LSEs.

**Fig. 3.**
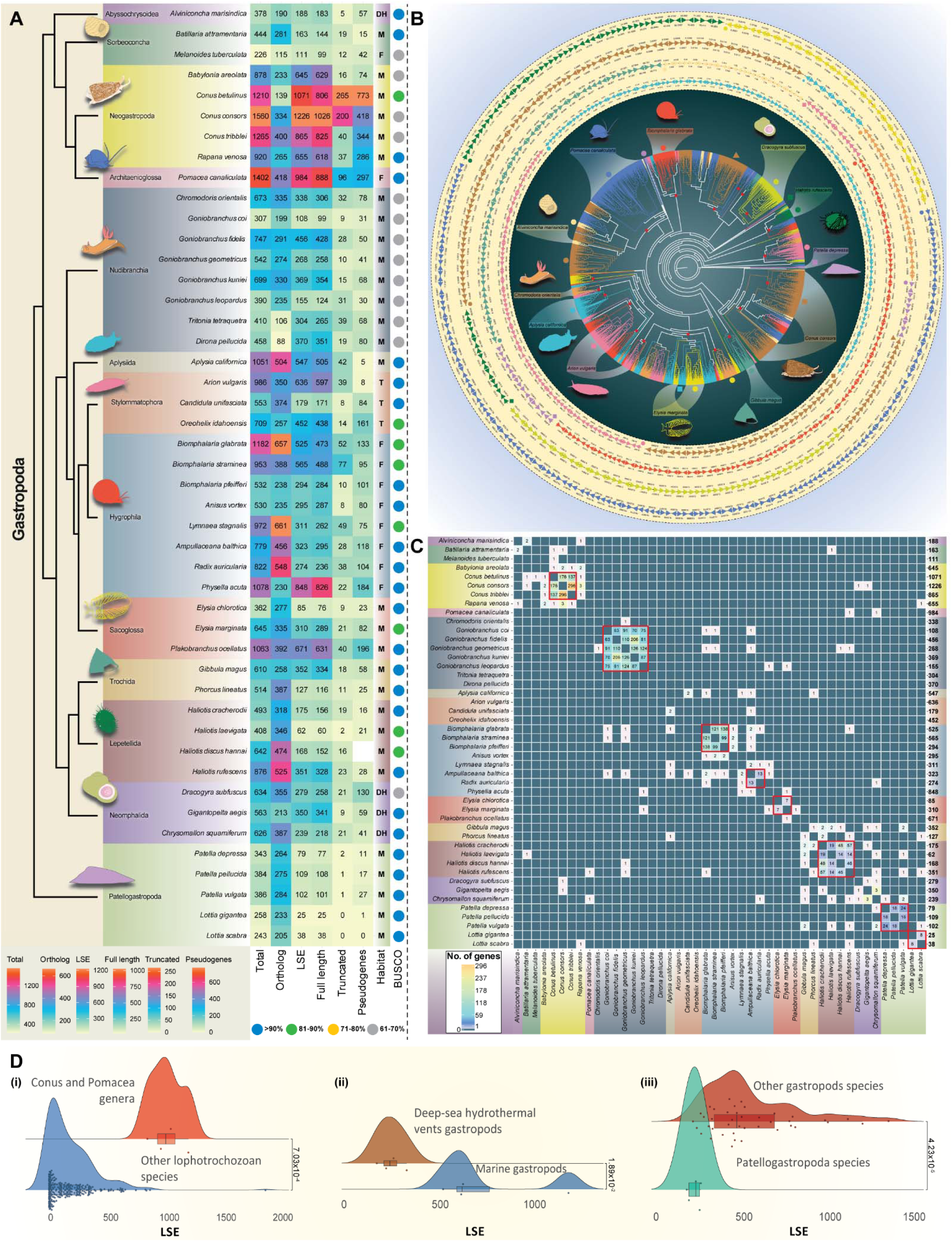
Comparative genomics and phylogenetic analysis of LSEs in Gastropods. **(A)** Phyletic representation of gastropods, showing taxonomic classifications, gene counts, and LSE classifications (full-length, truncated, pseudogene). Habitat distribution is represented as M (marine), F (freshwater), T (terrestrial), and DH (deep-sea hydrothermal vents). Colored dots indicate BUSCO genome completeness scores. **(B)** Maximum-likelihood (ML) tree displaying lineage-specific clustering of Class-A GPCRs from representative taxa across analyzed orders, with high-confidence supports (red circles, bootstrap values >90%). The outer layer shows the organization of LSE-forming genes on genomic scaffolds, with arrowheads denoting genes and numbers indicating gene locations in megabases. Colored shapes at scaffold beginnings correspond to tree topology clusters. **(C)** Heatmap illustrating the absence of one-to-one orthologs between species for classified LSEs using OrthoFinder inference. Cells with zero one-to-one orthologs are highlighted in teal. One-to-one orthologs are primarily found in intra-genus comparisons (red squares). Few orthologs were observed outside these comparisons in the OrthoFinder analysis, but they align within LSE-forming clusters in phylogenetic trees (see supplementary material). The number of LSE-forming genes used for cross-validation with OrthoFinder is shown on the right. Panel **(D)** presents statistical analyses. Significant differences in LSE counts: (i) between Conus and Pomacea genera compared to other lophotrochozoan species (W = 931) (left); (ii) between gastropods in deep-sea hydrothermal vents and marine environments (t = -3.19; df = 6) (center); (iii) between members of the Patellogastropoda order and other gastropods (W = 6) (right). The Wilcoxon rank sum exact test was used for (i) and (iii), and the two-sample t-test for (ii), with p < 0.05 considered significant.

**Fig. 4.**
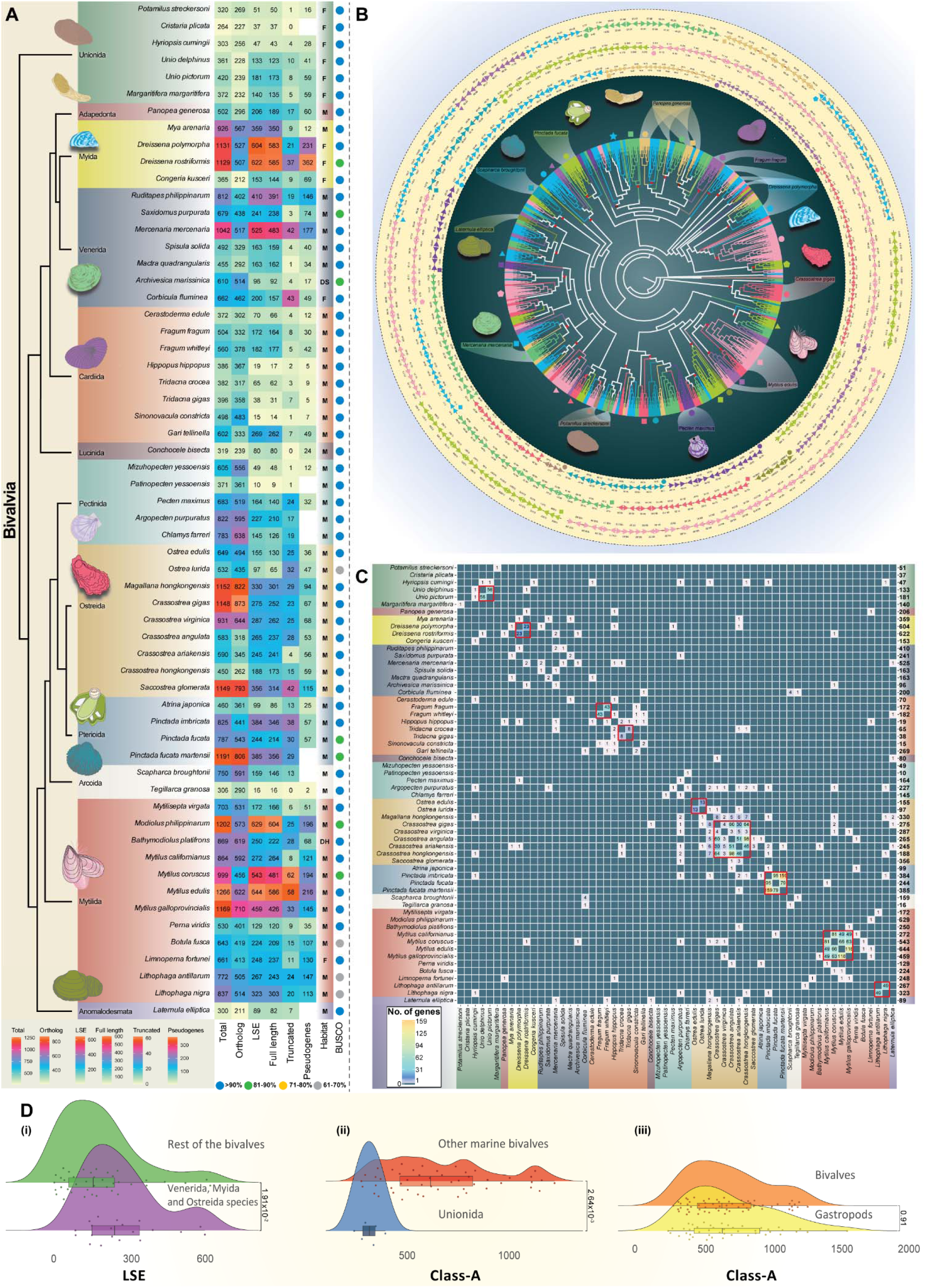
Phyletic distribution and comparative genomic analysis of LSEs in Bivalvia. **(A)** Phyletic representation of bivalves, detailing taxonomic classifications, gene counts, and LSE classifications (full-length, truncated, pseudogene). Habitat distribution: M (marine), F (freshwater), T (terrestrial), DS (deep-sea), and DH (deep-sea hydrothermal vents). Colored dots indicate BUSCO genome completeness scores. **(B)** ML tree displaying LSEs from representative taxa across analyzed bivalve orders, with high-confidence supports (red circles, bootstrap values >90%). The organization and illustration of scaffolds are as depicted in Fig.3. **(C)** Heatmap illustrating the absence of one-to-one orthologs between species for classified LSEs using OrthoFinder analysis. The number of LSE-forming genes used for cross-validation with OrthoFinder is shown on the right. The color scheme and the description of the OrthoFinder plot are consistent with Fig.3. Panel **(D)** presents statistical analyses: (i) significant variations in LSE counts among Venerida, Myida, and Ostreida compared to other bivalves (W = 550) (left); (ii) significant differences in class-A GPCR counts between Unionida freshwater mussels and other marine bivalves (center); (iii) no significant difference in class-A GPCRs between bivalves and gastropods (W = 1397.5) (right). Wilcoxon rank sum tests with continuity correction were applied for (i) and (iii), while (ii) utilized Kruskal-Wallis rank sum tests, followed by Dunn’s posthoc analysis with Bonferroni correction. Differences with p < 0.05 were considered statistically significant.

#### Sea Slugs with well-developed sensory structures encode large LSEs

Shell-less sea slugs, known for their vibrant colors and sluggish movement, possess well-developed sensory structures such as rhinophores and oral tentacles, with chemoreception serving as one of the principal mechanisms aiding in navigation, prey location, and foraging (*5, 27–30*). Consistent with this, the previously identified Class-A GPCR LSEs in *Aplysia californica*, specifically enriched in external sensory tissues, formed well-defined and distinct clusters in tree topologies, highlighting adaptations to their environmental niche (*24*). Our analysis, which incorporated 11 additional genomes of sea slugs from the Nudibranchia and Sacoglossa orders, reveals similar expansions forming distinct clusters with no identifiable orthologs compared to members from other orders (Class-A: 307-1063 genes; mean: 612; LSEs: 85-671; mean: 331; 12 species analyzed) (Fig. 3B and C, fig. S1A). The OrthoFinder analysis reinforces the absence of orthologous relationships between *Aplysia* LSEs (547 sequences) and those of *Chromodoris orientalis* (338) or taxa from the *Goniobranchus* and *Elysia* genera (Fig. 3C). However, intra-genus comparisons, exemplified by five genomes of the genus *Goniobranchus*, indicate a substantial increase in the proportion of orthologs (Fig. 3C), suggesting that lineage-specific diversifications are more prominent at the order and inter-family level comparisons. Despite an increase in the number of orthologs within the same genus, a notable proportion (around 24%) does not form orthologous groups, indicating considerable diversity even at the intra-genus level (Fig. 3C and fig. S2A). Similar occurrences were observed in various molluscan genera when multiple genomes were accessible for comparative analysis (see sections below).

#### Neogastropoda: the case of predatory mollusks

The members of Neogastropoda are recognized as predatory carnivorous mollusks, exhibiting varying degrees of predatory behavior, from actively seeking prey to grazing on sessile invertebrates and engaging in scavenging activities (*31–33*). Analysis of the genomes of three families, Conidae, Buccinidae, and Muricidae, revealed a significant increase in the proportion of Class-A GPCRs compared to mollusks from various other orders (P = 1.2x10^-3^, Wilcoxon rank sum test with continuity correction) (fig. S2B). Likewise, these families also show a high proportion of LSEs, comprising 68-88% of the Class-A GPCR repertoire (Class-A: 878-1560 genes; mean 1166; LSEs: 645-1226; mean 892; 5 species analyzed) (Fig. 3A-C, fig. S1A).

OrthoFinder analysis of these LSEs demonstrated similar trends (Fig. 3C), with no identifiable orthologous relationships observed at inter-family level comparisons. Interestingly, when comparing LSEs identified at the order level in three genomes of the genus *Conus*, there was only a modest increase, with just 19% of identified sequences displaying one-to-one orthologous relationships (Fig. 3C, fig S2A), highlighting significant diversity even within the genus.

Comparing the LSEs of Neogastropoda with related orders (Abyssochrysoidea, Architaenioglossa, Sorbeoconcha) within Caenogastropoda (*34*) revealed distinctive patterns. Specifically, the Class-A GPCR counts and their corresponding LSE components suggest independent and significant expansions in the stem lineage leading to Neogastropoda and Architaenioglossa (Fig. 3A). These expansions likely reflect adaptations to diverse habitats as species of the order Neogastropoda are primarily marine, found in coral reefs, rocky shores, seagrass beds, intertidal zones, and deep seas, while Architaenioglossa members inhabit terrestrial environments like meadows, grasslands, and forests (Fig. 3A and table S1).

#### Distinct expansions in amphibious freshwater and air-breathing land snails

While marine gastropods exhibit significant expansions, we investigated whether comparable trends exist in freshwater snails, of which some occasionally manifest amphibious traits (*35–38*). Analyzing 10 genomes across five diverse families across three distinct orders unveiled notable expansions, with LSEs constituting about 30-80% of the Class-A GPCR repertoire (Class-A: 226-1402 genes; mean: 847; LSEs: 111-984; mean: 453; 10 species analyzed) (Fig. 3A-C). In particular, *Pomacea canaliculata* (984), *Physella acuta* (848), and *Biomphalaria glabrata* (525) displayed significant LSE proportions, lacking apparent orthologous relationships when compared to each other (Fig. 3A-C, fig. S1A). Limiting to family-level comparisons among freshwater snails also revealed a considerable degree of diversification. Comparing three distinct genomes of the *Biomphalaria* genus with *Anisus vortex* from the family Planorbidae revealed minimal orthologous relationships, with fewer than five genes displaying one-to-one orthologous associations in distinct pairwise comparisons (Fig. 3C and fig S2A). Similar trends were observed for members of the family Lymnaeidae, and LSEs were distinctly different at the level of inter-family comparisons within the same and across orders (Fig. 3A-C). These findings suggest that despite the common occurrence of freshwater snails in ponds, lakes, rivers, and marshes, the observed LSEs indicate diversifications likely influenced by distinct environmental factors and conditions specific to each habitat (*35*). Similar observations were noted when examining terrestrial air-breathing snails such as *Arion vulgaris* (636) and *Oreohelix idahoensis* (452) and other members from three distinct families of the order *Stylommatophora* (Fig. 3A). These observations align with their habitat diversity, with *Arion vulgaris* typically found in moist and humid environments, *Candidula unifasciata* thriving in dry and sunny lowlands, and *Oreohelix idahoensis* inhabiting montane and subalpine areas (Fig. 3A).

#### LSEs in both marine and freshwater bivalves

Of the 12 analyzed bivalve orders, taxa from 11 different predominantly inhabit diverse marine environments, including intertidal zones, shallow coastal waters, estuarine areas, and sandy or muddy substrates (Fig. 4A and table S1). Particularly notable are the significant expansions observed in the Venerida, Myida, and Ostreida orders compared to other bivalves (Fig. 4A-D, fig. S1B). This is illustrated by taxa such as *Mercenaria mercenaria* (525), *Dreissena rostriformis* (622), *Dreissena polymorpha* (604), and *Crassostrea angulata* (265), constituting 45-55% of their Class-A GPCRs (Class-A: 365-1152 genes; mean: 774; LSEs: 96-622; mean: 286; 20 species analysed; P = 1.91x10^-2^, Wilcoxon rank sum test with continuity correction) (Fig. 4D). Similar to gastropods, the lack of apparent orthologous relationships is evident in inter-order, inter-, and intra-family level comparisons of LSEs in bivalves, with only a slight increase in intra-genus comparisons (Fig. 4C). An analysis of four genomes from the *Mytilus* and *Crassostrea* genera reveals that approximately 22% and 28% of the LSEs derived from family and order-level comparisons establish one-to-one orthologous relationships when confined to the intra-genus level (Fig. 4C). Similarly, examinations of three genomes within the *Pinctada* genus show that 32% of genes form orthologous relationships, while two genomes from each of the *Fragum* and *Tridacna* genera exhibit only 12% and 8%, respectively (Fig. 4C).

Unlike most marine bivalves with an average of over 690 Class-A GPCRs, freshwater mussels of the Unionida order show a significant decrease ranging from 264 to 420 receptors (P = 2.64x10^-3^, Kruskal-Wallis rank sum test followed by Dunn’s post-hoc test with Bonferroni correction) (Fig. 4D). However, freshwater mussels from the Dreissenidae family of the order Myidae, such as *Dreissena polymorpha* and *Dreissena rostriformis*, have Class-A GPCR numbers similar to marine bivalves without significant difference, each surpassing 1100 GPCRs (P = 0.34, Kruskal-Wallis rank sum test followed by Dunn’s post-hoc test with Bonferroni correction). Likewise, approximately 54% of these form LSEs, and no decrease is observed when limiting the comparisons solely to the intra-genus level (Fig. 4C). Together these observations suggest high levels of interspecific variations.

#### Relative reduction in abyssal gastropods, bivalves, and lineages with specific adaptations

Next, we investigated whether there are discernible patterns among gastropods and bivalves that have adapted to deep-sea hydrothermal vents and other specific conditions. Markedly, *Gigantopelta aegis*, *Chrysomallon squamiferum*, *Dracogyra subfuscus*, and *Alviniconcha marisindica* show significant reductions in Class-A GPCRs and corresponding LSEs (Class-A: 378-634 genes; mean: 550; LSEs: 188-350; mean: 264; 4 species analyzed) (Fig. 3A). While these are sizeable counts, it is a considerable decrease compared to other marine gastropods like Neogastropods and Aplysiidae which typically encode a much higher proportion, or twice as many as LSEs (Class-A: 307-1560 genes; mean: 816; LSEs: 108-1226; mean: 491; 22 species analyzed; Class-A: P = 1.67x10^-2^, two-sample t-test; LSEs: P = 1.89x10^-2^, two-sample t-test) (Fig. 3D and fig. S2B). This relative reduction is also in congruence with recent investigations into opsin gene evolution, suggesting a significant gene loss in *Gigantopelta aegis* (*39, 40*). This might emphasize that adaptation to these extreme conditions likely drives contractions in chemosensory gene families, while on the contrary, expansions have occurred in gene families linked to metabolism, DNA stability, antioxidation, and biomineralization, indicating varied selection pressures in shaping the evolution of these gene families in extreme environments (*39, 40*). Similarly, within the bivalve category, *Archivesica marissinica* of the Vesicomyidae family and *Conchocele bisecta* of the Thyasiridae family, residing in deep-sea hydrothermal and methane-rich vents, also exhibit reductions with only 96 and 80 receptors, forming the LSE component (Fig. 4A). Besides deep-sea mollusks, prominent reductions were also observed in limpets belonging to the Patellogastropoda order, with genomic analysis of the Patellidae and Lottiidae families showing decreases in both Class-A GPCRs and the corresponding LSE component compared to other gastropods (Class-A: P = 2.87x10^-4^, Wilcoxon rank sum exact test; LSEs: P = 4.23x10^-5^, Wilcoxon rank sum exact test) (Fig. 3A and D, fig. S2B). While the cause of this reduction in limpets is unclear, their sedentary lifestyle, attaching firmly to specific locations for extended periods and grazing on algae and microorganisms on rocky surfaces using their muscular foot, may explain the observed decrease (*41*). Similarly, bivalves like *Congeria kusceri* (Dreissenidae), which are adapted to subterranean waters with a highly confined biogeographical distribution, exhibit notable reductions compared to other Dreissena species. They are characterized by limited sensory organs and a sedentary lifestyle, attaching to cave walls (*42*).

#### Decreased proportions of LSEs across all examined cephalopod orders

Similar to gastropods and bivalves, cephalopods display a wide distribution across diverse habitats, from pelagic to benthic environments, including various coastal settings, highlighting their versatility and adaptation to diverse ecological niches (*43, 44*). However, from a comparative perspective to gastropods and bivalves, examining 19 genomes across nine families and four orders revealed a notable pattern of a consistent reduction in both Class-A GPCR count and the LSE component across all analyzed cephalopods (Class-A: 46-411 genes; mean: 183; LSEs: 3-146; mean: 29; 19 species analysed; Class-A: P < 2.2x10^-16^, Wilcoxon rank sum test with continuity correction; LSEs: P < 2.2x10^-16^, Wilcoxon rank sum test with continuity correction) (Fig. 5A-D and 5I, fig. S1C). Another observation within cephalopods is the significant reduction observed in six specific taxa: *Octopoteuthis deletron*, *Japetella diaphana*, *Architeuthis dux*, *Argonauta argo*, *Muusoctopus leioderma,* and *Muusoctopus longibrachus* belonging to five distinct families that have independently adapted to deep-sea environments (*45*). This parallels the findings in abyssal gastropods and bivalves (Class-A: 46-323 genes; mean: 141; LSEs: 6-34; mean: 17; 3 species analysed) (P = 4.79x10^-2^, Wilcoxon rank sum test with continuity correction) (Fig. 5I).

**Fig. 5.**
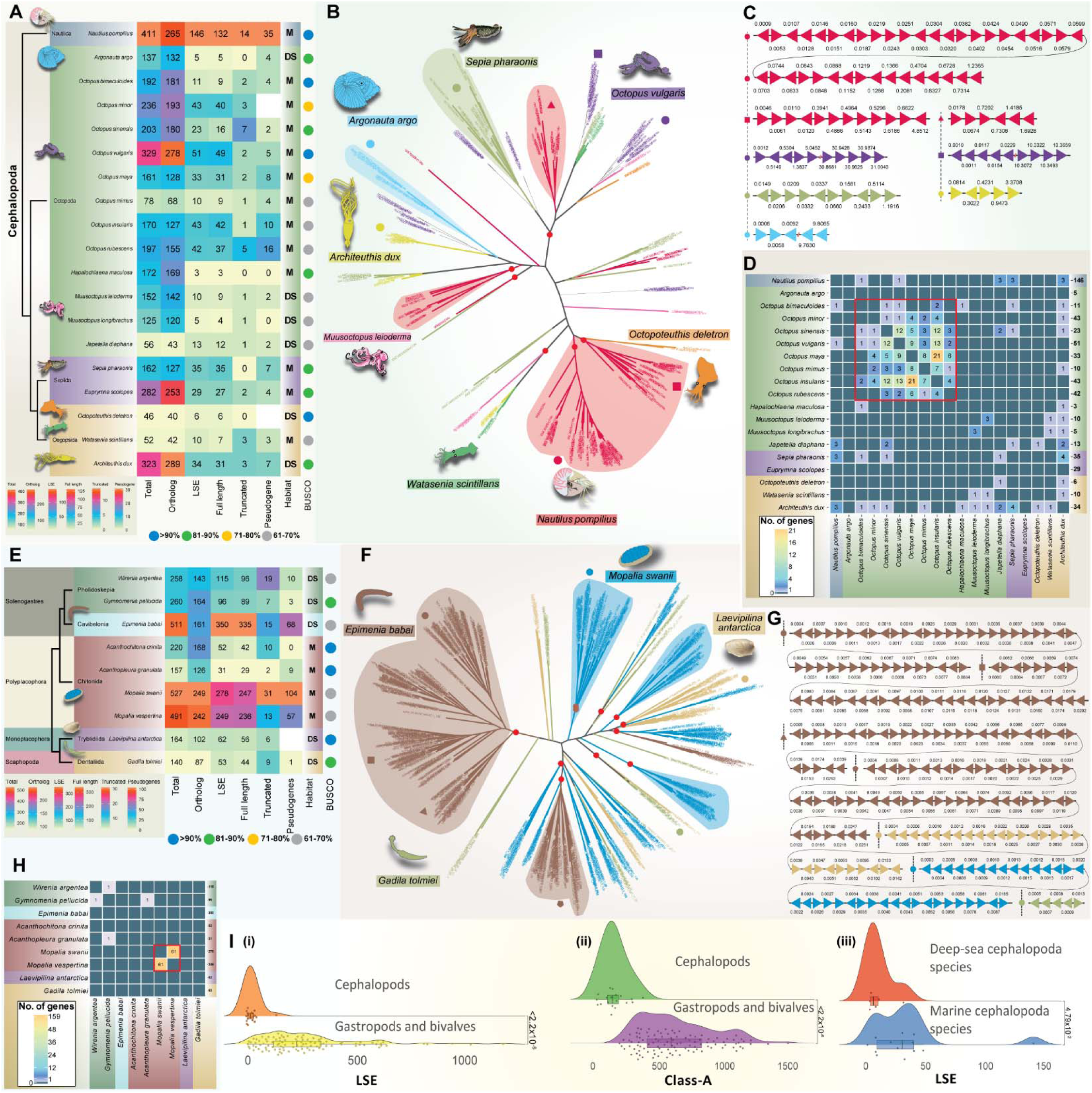
Distribution and observed dynamics of LSEs in cephalopods and other molluscan classes. **(A)** Phyletic representation of analyzed cephalopods, including taxonomic classifications, gene counts, and classifications of LSEs (full-length, truncated, pseudogene). Habitat distribution: M (marine) and DS (deep-sea). The heatmap color scheme and description are as in Figs. 3 and 4. **(B)** ML tree displaying LSEs from representative cephalopod taxa with high-confidence support (bootstrap values >90% indicated by red circles). **(C)** Schematic representation of clustering of LSE genes in genomic scaffolds. (D) Heatmap illustrating the absence of one-to-one orthologs between species for classified LSEs using OrthoFinder analysis. The color scheme and description of the plot are consistent with Figs. 3 and 4. Panels **(E-H)** depict comparative analysis and observed patterns in other molluscan classes. (E) Phylogenetic distribution showing gene counts, classifications, and habitat distributions (M for marine, DS for deep-sea). (F) ML tree topology displaying LSE clustering. (G) Schematic representation of clustering of LSE genes in genomic scaffolds. (H) OrthoFinder plots. The color scheme, descriptions, and layouts are detailed similarly to Figs. 3 and 4. Panels **(I)** show statistical analyses: significant differences in LSEs (W = 77) and class-A GPCRs (W = 0) between cephalopods and combined gastropods/bivalves (i and ii); (iii) significant differences in LSE counts between deep-sea and marine cephalopod species (W = 16). Wilcoxon rank sum test with continuity correction was used for all three plots; differences with p < 0.05 were considered significant.

Intriguingly, the *Nautilus pompilius*, representing the earliest branching lineage among all cephalopods (*46*), has the highest count of Class-A receptors, specifically 411, with an LSE component consisting of 146 receptors (Fig. 5A). This may indicate a slight reduction in the lineage leading to coleoid cephalopods and the availability of additional genomes within the order Nautilida would further clarify this. Overall, the intriguing decrease in Class-A GPCR LSEs across most cephalopods raises questions, especially considering their advanced sensory systems. A potential explanation could lie in the existence of alternative chemosensory gene families, like cephalopod-specific chemotactile receptors, where ionotropic receptors play a vital role in sensing environmental cues within their ecological niche (*47–49*). Additionally, the ability of octopod arm suckers, decapod buccal lips, and nautiloid tentacles to detect molecules through direct contact, facilitated by the presence of gustatory receptors, further adds complexity to the sensory landscape, and could potentially argue for the relative reduction in GPCR mediated chemoreception compared to gastropods and bivalves (*47–50*).

#### Patterns in other molluscan classes: from Chitons to Tusk Shells

We also investigated other molluscan classes with limited genomic data: Polyplacophora, Monoplacophora, Solenogastres, and Scaphopoda. Chitons from Polyplacophora inhabit diverse marine environments, from intertidal zones to deep-sea habitats. Monoplacophora, known for their primitive nature, dwell in ocean depths. Solenogastres occupy various marine habitats, including hydrothermal vents and intertidal regions, while Scaphopoda, or tusk shells, burrow into sandy or muddy substrates for feeding and respiration. Within Polyplacophora, Mopaliidae genomes exhibit nearly 500 Class-A GPCRs, with over half being LSEs; Acanthochitonidae and Chitonidae members have about 200 Class-A GPCRs, with fewer than 30% classified as LSEs (Fig. 5E-H, fig. S1D). Intriguingly, the deep-sea-adapted Solenogastres exhibit considerable expansions in taxa belonging to both Gymnomeniidae and Epimeniidae families. For example, *Epimenia babai* encodes 511 Class-A GPCRs, of which 350 are LSEs. Conversely, the deep-sea Monoplacophora and Scaphopoda classes encode only 160 Class-A GPCRs. These findings underscore diverse patterns, and the availability of additional genomes within each family and class will confirm whether these are consistent trends.

### Observed Trends Across Annelid Diversity

With a global distribution, members of the phylum Annelida exhibit diverse habitats, ranging from marine, terrestrial, freshwater, and deep-sea hydrothermal vents (*51–53*). Similar to Mollusks, we witness a range of trends within different orders inhabiting distinct ecological niches, showcasing both expansions and reductions.

#### Expansions in both marine and terrestrial annelids

An analysis of 20 genomes spanning 12 families of Polychaeta worms and an additional genome from the Sipunculidae family inhabiting the diverse marine environments reveals notable expansions (Fig. 6A-C, fig. S1E) (Class-A: 191-1481 genes; mean: 645; LSEs: 88-889; mean: 335; 21 species analyzed; P = 2.8x10^-4^, Wilcoxon rank sum test with continuity correction) (Fig. 6D). Particularly, species like *Capitella teleta* (1481 Class-A genes; 857 LSEs) and *Harmothoe impar* (1264 Class-A genes; 889 LSEs) exhibit large LSEs (Fig. 6A-C, fig. S1E). Similarly, within the Clitellata class of terrestrial annelids, all five genomes from three different families of the order Crassiclitellata exhibit sizable Class-A GPCR counts and corresponding LSE. Moreover, these identified LSEs in both marine and terrestrial annelids do not show an increase in one-to-one orthologous relationships at the intra-family level comparisons (Fig. 6C). It would be intriguing to observe if similar trends emerge at the intra-genus level with more genomes available for comparison.

**Fig. 6.**
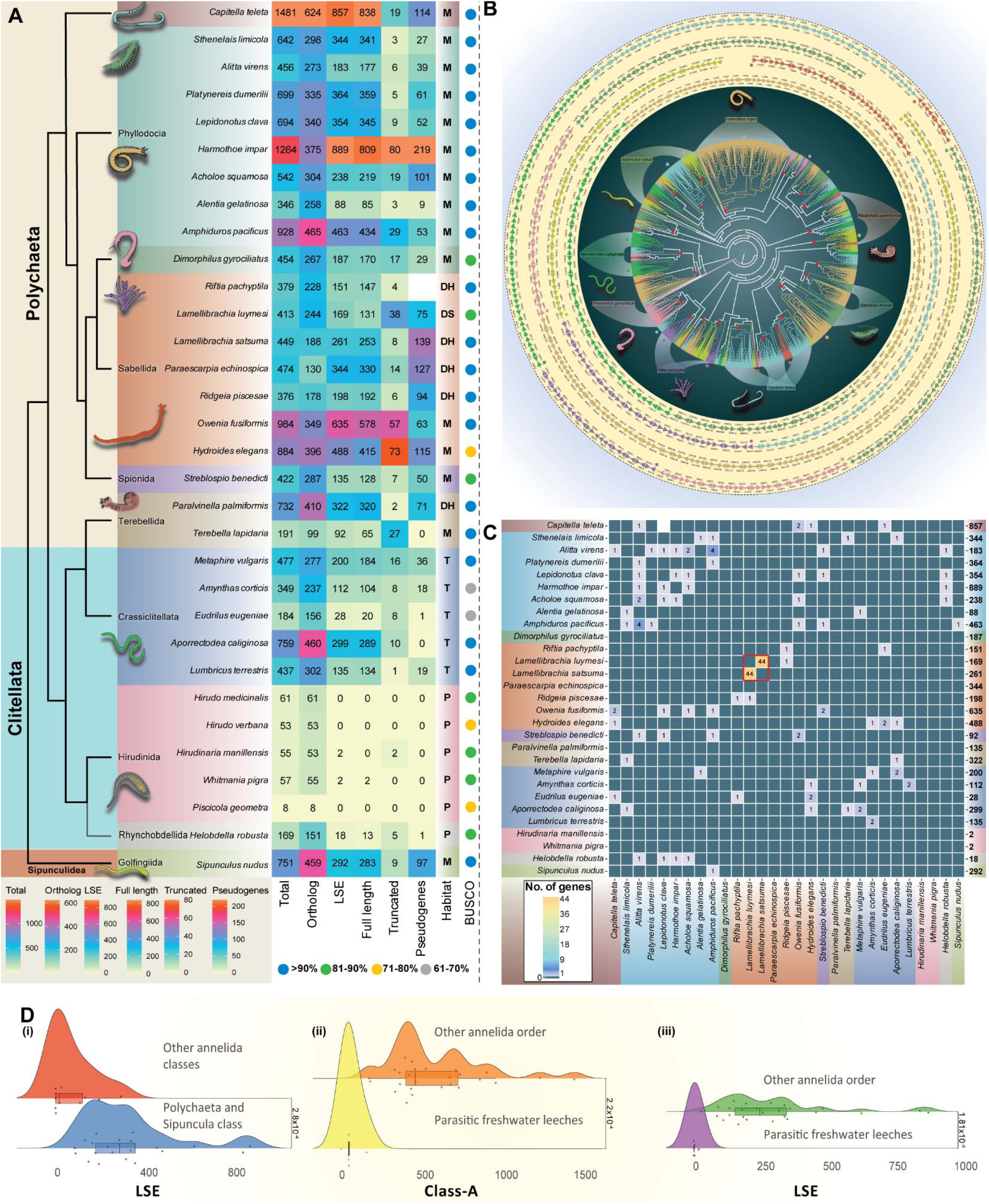
Comparative analysis and evolutionary dynamics of LSEs in Annelids. **(A)** Phyletic representation of analyzed annelids, detailing taxonomic classifications, gene counts, and LSE classifications (full-length, truncated, pseudogene). Habitat distribution: M (marine), T (terrestrial), P (parasitic), DS (deep-sea), and DH (deep-sea hydrothermal vents). Colored dots indicate BUSCO genome completeness scores. **(B)** ML tree displaying LSEs from representative annelids with high-confidence support (bootstrap values >90% indicated by red circles). The organization and illustration of scaffolds are as depicted in Figs. 3 and 4. **(C)** Heatmap illustrating the absence of one-to-one orthologs between species for classified LSEs using OrthoFinder analysis. The color scheme and description of the plot are consistent with Figs. 3-5. Panels **(D)** presents statistical analyses: (i) significant differences in LSEs between class Polychaeta and Sipunculus and the rest of the Annelida classes (W = 207.5); (ii and iii) between parasitic freshwater leeches of the order Hirudinida and Rhynchobdellida and rest of the Annelida species in both class-A GPCRs (t = -4.2, df = 30) and LSE (W = 0), respectively. Wilcoxon rank sum test with continuity correction was used for (i) and (iii), and a two-sample t-test for equal variances for (ii), with p < 0.05 considered significant.

#### Reductions among the parasitic freshwater leeches

Examining five genomes from three distinct families within the Hirudinida order and one genome from the Rhynchobdellida order reveals a significant reduction in the number of Class-A GPCRs and their corresponding LSEs across all studied parasitic freshwater leeches (Class-A: 8-169 genes; mean: 67; LSEs: 0-18; mean: 3.6; 6 species analyzed; Class-A: P = 2.2x10^-4^, two-sample t-test; LSEs: P = 1.81x10^-4^, Wilcoxon rank sum test with continuity correction) (Fig. 6A-D). For instance, in the Hirudinidae family, the number of Class-A GPCRs in the three analyzed genomes ranges from 53 to 61 receptors with no prominent LSEs (Fig. 6A). Despite the expected role of chemoreception in prey detection, the sensory mechanisms of leeches remain largely unclear. Reports indicate that mechanosensation and photosensory inputs might be the primary methods for environmental sensing, which could explain the significant reduction in GPCR-mediated chemoreception (*51, 54–57*). Our findings highlight the need for further investigation into the chemosensory behavior mechanisms of these parasitic leeches.

#### The case of deep-sea annelids

In contrast to abyssal gastropods and bivalves, an analysis of five genomes from hydrothermal vent worms (Siboglinidae family, order Sabellida) reveals a relatively sizable repertoire of Class-A GPCRs and corresponding LSEs (Class-A: 376-474 genes; mean: 418; LSEs: 151-344; mean: 224; 5 species analyzed) (Fig. 6A-C). These counts slightly exceed those typically found in most deep-sea gastropods and bivalves and are comparable to deep-sea-adapted Solenogastres (supported by no significant statistical difference between Solenogastres and deep-sea annelids: P = 0.94). However, no significant statistical difference was also found between deep-sea annelids and deep-sea gastropods/bivalves in general (P = 0.85), suggesting the need for more Annelida genomes to clarify this further.

### Evolutionary Shifts in Platyhelminthes: From Expansions in Free-Living Ancestors to Reductions in Parasitic Forms

Platyhelminthes show diverse forms, from free-living in aquatic and terrestrial environments to parasitic species with complex life cycles involving various hosts (Fig. 7A and table S1) (*58, 59*). Despite initial classification outside Lophotrochozoa, recent analyses consistently group them within this category (*60*).

**Fig. 7.**
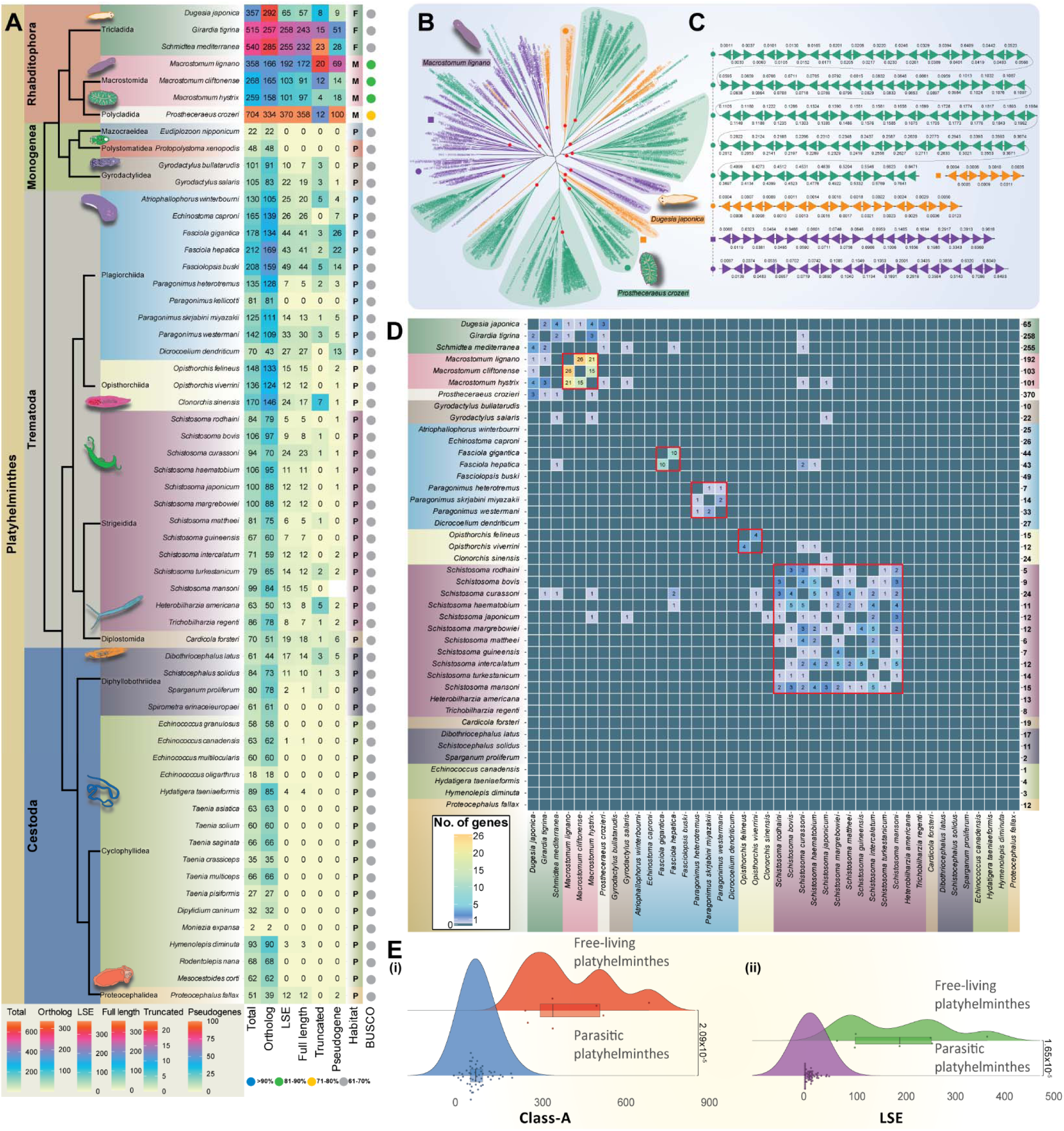
Comparative genomic analysis informs contrasting patterns between free-living and parasitic Platyhelminthes. **(A)** Phyletic representation of analyzed Platyhelminthes, detailing taxonomic classifications, gene counts, and LSE classifications (full-length, truncated, pseudogene). Habitat distribution: F (freshwater), M (marine), and P (parasitic). Colored dots indicate BUSCO genome completeness scores. **(B)** ML tree displaying LSEs from representative Platyhelminthes with high-confidence support (bootstrap values >90% indicated by red circles). **(C)** Schematic representation of clustering of LSE genes in genomic scaffolds. **(D)** Heatmap illustrating the absence of one-to-one orthologs between species for classified LSEs using OrthoFinder inference. The color scheme and description of the plot are consistent with Figs. 3-6. (**E)** Statistical analysis shows significant differences in (i) class-A GPCR counts (W = 364) and (ii) LSE counts (W = 0) between parasitic and free-living Platyhelminthes. The Wilcoxon rank sum test with continuity correction was used, with p < 0.05 considered significant.

Examining 52 genomes across ten orders of parasitic Platyhelminthes—Trematoda (Flatworms), Cestoda (Tapeworms), and Monogenea (Roundworms)—reveals a marked reduction in Class-A counts, with few or no LSEs compared to free-living Platyhelminthes (Class-A: 2-212 genes; mean: 88; LSEs: 0-49; mean: 10; 52 species analyzed; Class-A: P = 2.09x10^-5^, Wilcoxon rank sum test with continuity correction; LSEs: P = 1.65x10^-5^, Wilcoxon rank sum test with continuity correction) (Fig. 7A-E, fig. S1F). While exceptions exist, such as members of the Fasciolidae family within Trematoda exhibiting a range of 178-212 GPCRs, with LSEs ranging from 43-49, most examined genomes contain fewer than 100 Class-A GPCRs. For instance, *Schistosoma mansoni*, a well-known parasite causing schistosomiasis, encodes approximately 99 Class-A GPCRs (*61*). Platyhelminthes, with distinct parasitic modes—Monogenea being ectoparasitic and Cestoda/Trematoda being endoparasitic—have evolved various adaptations (*58, 62*), including tegument formation, specialized attachment structures, body elongation, complex life cycles, and reductions in sensory and locomotor structures compared to free-living ancestors (*63, 64*). At the genomic level, adaptations to parasitism also marked a substantial reduction in homeodomain-containing genes, piwi, vasa, Wnt, NEK kinases, fatty acid biosynthesis, and opsin genes (*39, 65, 66*). Our survey further validates these patterns, uncovering a significant decrease in GPCR-mediated chemoreception in parasitic Platyhelminthes.

In contrast, both freshwater and marine free-living Platyhelminthes exhibit noticeable expansions, as evidenced by the analysis of seven genomes across three distinct orders (Class-A: 259-704 genes; mean: 428; LSEs: 65-370; mean: 192; 7 species analyzed). Specifically, *Prostheceraeus crozeri*, a member of the Euryleptidae family, possesses around 704 Class-A GPCRs, with 370 as LSEs (Fig. 7A-D, fig. S1F). This augmentation suggests a prevalence of GPCR-mediated chemoreception in these free-living Platyhelminthes, which in general aligns with the presence of sensory structures such as ciliated epidermis, rhabdites, statocysts, and tentacles or auricles (*67, 68*). Thus, the ancestral free-living Platyhelminthes possess LSEs comparable to other free-living lophotrochozoans, with significant reductions observed in lineages leading to parasitic forms akin to patterns in parasitic leeches.

### Trends in other Lophotrochozoan Phyla

We also explored various additional phyla within the Lophotrochozoa, including Bryozoa, Phoronida, Entoprocta, Nemertea, and Brachiopoda that inhabit a wide array of marine environments (Fig. 8A). Significant expansions in Class-A and corresponding LSE counts were observed in Nemertea, compared to Entoprocta, Bryozoa, Phoronida and Brachiopoda (Class-A: P = 2.99x10^-6^, Wilcoxon rank sum test with continuity correction; LSEs: P = 5.65x10^-7^, Wilcoxon rank sum test with continuity correction) (Fig. 8A-D, fig. S1G). For example, *Notospermus geniculatus* exhibits approximately 2352 Class-A GPCRs, with 1904 as LSEs. *Lineus longissimus*, another Lineidae member, encodes approximately 1009 Class-A GPCRs, with 764 as LSEs. Though the reasons for their large expansions are unclear, these worms have advanced sensory systems and display predatory and scavenging behaviors in marine environments, using an eversible proboscis for hunting and protecting themselves with a neurotoxin-infused mucus layer (*69–71*). Studies suggest an expansion of toxin and mucus-secreting genes in *Notospermus geniculatus*, possibly as adaptations to a predatory lifestyle (*72*). Further exploration of Nemertea genomes will determine if gene expansions are common or if contractions occur in certain classes. Expansions were also found in all analyzed Brachiopoda and Phoronida members (Fig. 8A-D, fig. S1G). In contrast, Bryozoa have fewer Class-A GPCRs (Class-A: 153-198 genes; mean: 175; LSEs: 24-99; mean: 75; 5 species analyzed). Limited genomic data for these phyla highlights the need for future research to confirm these trends.

**Fig. 8.**
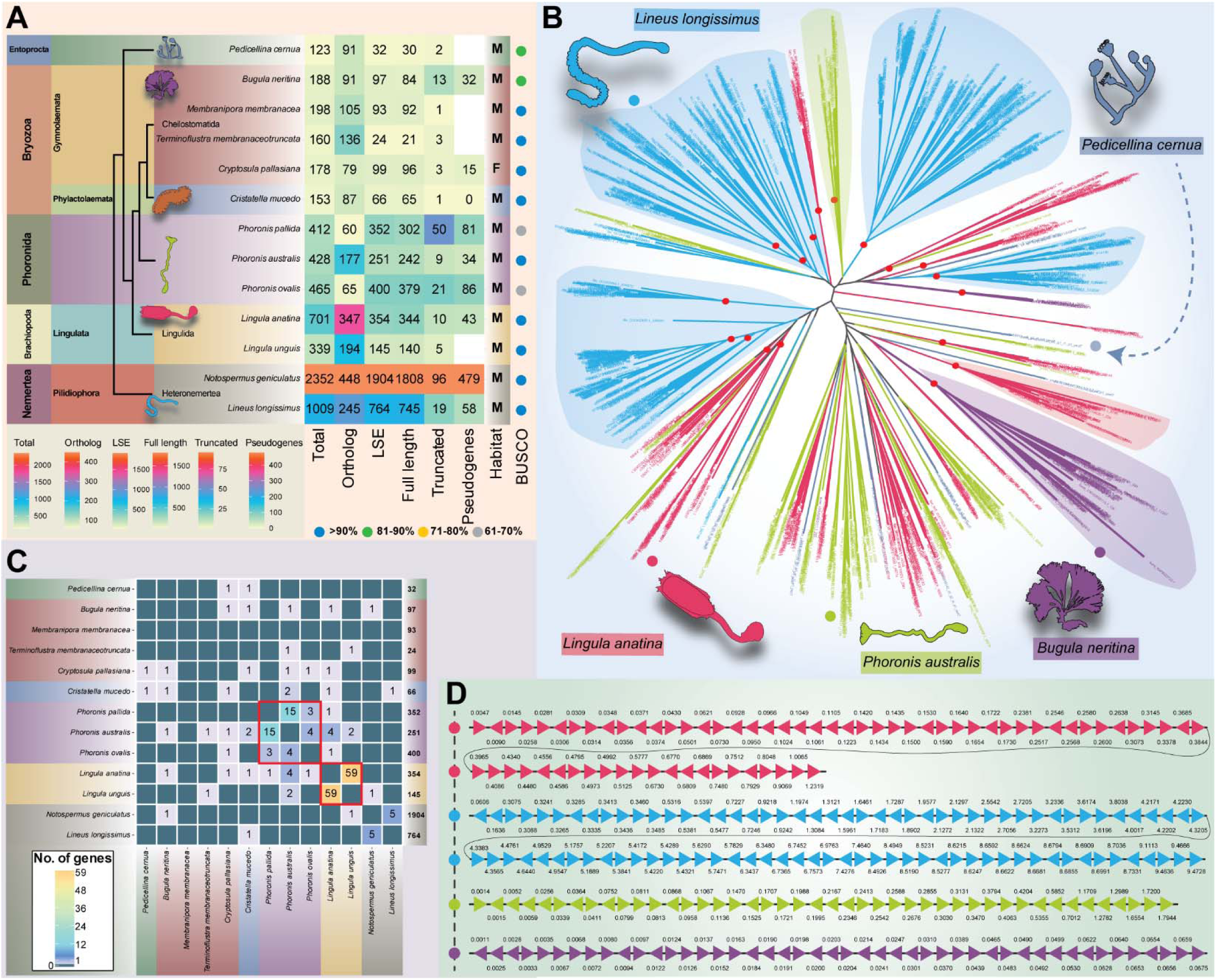
Phyletic distribution and overall trends in other lophotrochozoan phyla. **(A)** Phyletic representation of other analyzed lophotrochozoan phyla, detailing taxonomic classifications, gene counts, and LSE classifications (full-length, truncated, pseudogene). Habitat distribution: F (freshwater), M (marine). The heatmap color scheme and description are as in Figs. 3-7. **(B)** ML tree displaying LSEs from representative species with high-confidence support (bootstrap values >90% indicated by red circles). **(C)** Heatmap illustrating the absence of one-to-one orthologs between species for classified LSEs using OrthoFinder inference. The color scheme and description of the plot are consistent with Figs. 3-7. **(D)** Schematic representation of clustering of LSE genes in genomic scaffolds.

### Roles of WGDs in the Evolution of LSEs Inform Non-Uniformity and Selective Effect

Given the extensive LSEs, we explored whether these were predominantly driven by any observed whole-genome duplication (WGD) events. A thorough examination of the available literature (table S1) indicates that WGD events are highly sporadic across Lophotrochozoa. In Gastropoda, WGDs occurred at key points: (i) at the common ancestor of Stylommatophora, and (ii) within specific Neogastropoda families or potentially at their common ancestor (*73–77*). In Bivalvia, such events are noted in families like Ostreidae and Mytilidae (*77–79*) and uniquely at the base of Decapodiformes within Cephalopoda (*77*). Outside Mollusca, potential WGDs occurred in certain Annelida families (Megascolecidae, Glossiphoniidae) and at the base of the Macrostomida order (*80–82*), which includes free-living Platyhelminthes (table S1). When analyzing these events alongside the observed expansions of Class-A GPCRs, we found no consistent correlation between WGD events and significant expansions. While WGD events may have contributed to increased LSEs, particularly in lineages like Neogastropoda and free-living Platyhelminthes, this pattern does not uniformly extend to other lineages that have experienced WGD. For example, *Candidula unifasciata* in the Stylommatophora order shows a reduced GPCR repertoire (32% LSEs), contrasting with species like *Arion vulgaris* and *Oreohelix idahoensis* (63-64% LSEs), which do not exhibit any evidence for WGDs. Remarkably, among the 238 analyzed taxa, 52 taxa encode over 50% of the Class-A GPCR repertoire as LSEs, with only six of these taxa showing evidence of WGD events (table S1). No significant difference was observed in the percentage of LSEs between the 46 non-WGD taxa and the 6 WGD taxa (P = 0.3, Wilcoxon rank sum exact test). This suggests that the primary influence on the shaping of Class-A GPCR LSEs appears to be driven by environmental adaptations and the utilization of these receptors for chemosensory roles rather than solely the consequence of WGDs.

### Signatures and the Rate of Evolution of the Ligand-Binding Regions

Next, we explored the diversity in ligand-binding pockets of all identified LSEs using a structure-guided approach (See methods). The analysis shows that TM regions have moderate to high sequence heterogeneity, even within the same taxon, with extensive variations at higher taxonomic levels (Fig. 9). By extracting majority rule consensus sequences from each LSE cluster, we compared evolutionary rates using Shannon entropy (H) (*83, 84*) (Fig. 9). Key observations are: (i) Extracellular segments of each TM region exhibit significantly higher positional entropy, compared to the rest of the TM-segment, a consistent trend across all seven TM helices and taxonomic levels in lophotrochozoa (Fig. 9). (ii) Besides the common Class-A GPCR motifs like DRY and NPxxY, no conserved sequence motifs specific to habitats or taxonomic groups were found, indicating evolution through extensive positive selection. This high variability and fast evolution, reflected in higher positional entropies, explain the lack of conserved or habitat-specific signatures in the ligand-binding pockets of these LSEs. (iii) Similarly, the extracellular loops of these LSEs display high sequence heterogeneity, typically containing charged and polar residues. In terms of length, the patterns mirror those seen in vertebrate olfactory receptors: ECL2 is notably longer, spanning 25 to 30 residues (*85*), whereas ECL1 and ECL3 range from 15 to 20 residues across various groups (*86, 87*).

**Fig. 9.**
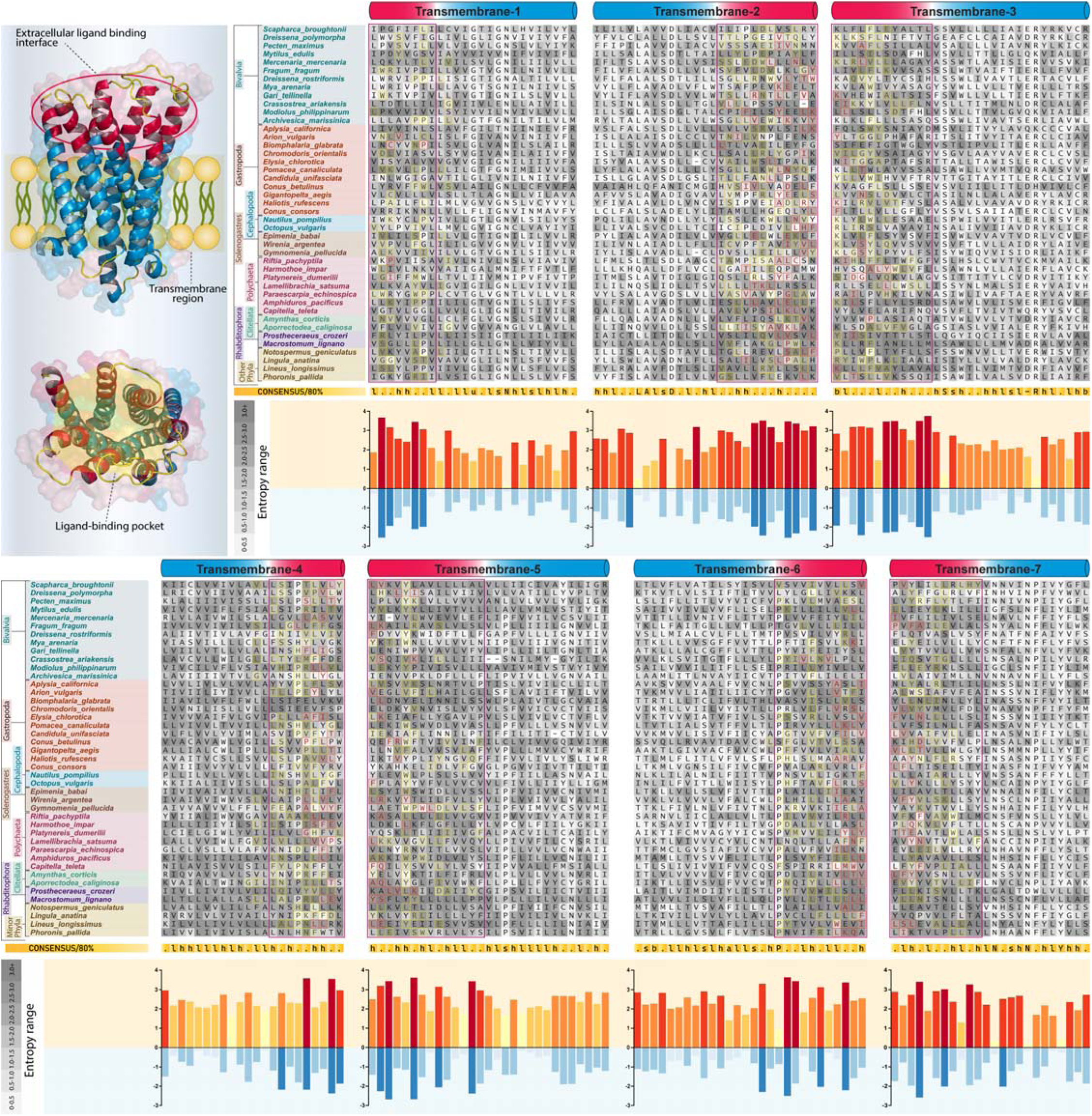
Multiple alignments illustrate the rate of evolution and signatures of positive selection in ligand-binding pockets of LSEs across phyla. Each sequence in the alignment is a consensus derived from the largest LSE cluster of the representative taxa. The background displays the entropy value of each consensus amino acid position from the corresponding LSE alignment, with varying shades of grey. Amino acids involved in inter-TM contacts are highlighted in yellow, and those interacting with the extracellular loop(s) are highlighted in red. Below each TM segment, a bar plot shows the overall entropy for the corresponding alignment position, with yellow to red indicating absolute entropy and light blue to dark blue indicating amino acid property-based entropy. The alignment’s topology is illustrated above, highlighting the extracellular ligand-binding regions of the alignment in red. High entropy regions at the ligand-binding interface are boxed in red in the alignment and bar plots. The left panel shows side and top views of a 3D class-A GPCR model highlighting the ligand-binding pockets.

We expanded our ligand-binding pocket analysis by focusing on residue contact networks at the apex of the 7TM helix bundle. Using AF2-predicted 3D structures from consensus sequences of LSE clusters, we analyzed 169 static structures across major phyla. Residue contact maps were generated using GetContacts, focusing on inter-TM and TM-ECL pairs, and the type of contacts between each residue pair was categorized based on their amino acid properties (see methods). In the analysis of inter-TM segments, variability in residue contact pairs and interaction types is evident across major phyla (Fig. 10A and B, and table S2). For instance, comparing 41 gastropod structures to 47 bivalve structures, gastropods show 511 unique residue contact pairs in 28 categories, whereas bivalves display 664 pairs in 30 categories. Annelids exhibit 495 pairs across 33 categories, contrasting with cephalopods’ 75 pairs in 14 categories. Overall, there are 35 categories of residue pairs across all phyla, with only 11 common to all, indicating diverse interaction networks. Notably, gastropods lack interactions in 7 categories, while cephalopods lack 21 of the 35 categories (table S2). While these numbers will vary with an increased sample size of analyzed structures, it is noteworthy that the variability observed in the alignments is reflected in the residue contact pairs.

**Fig. 10.**
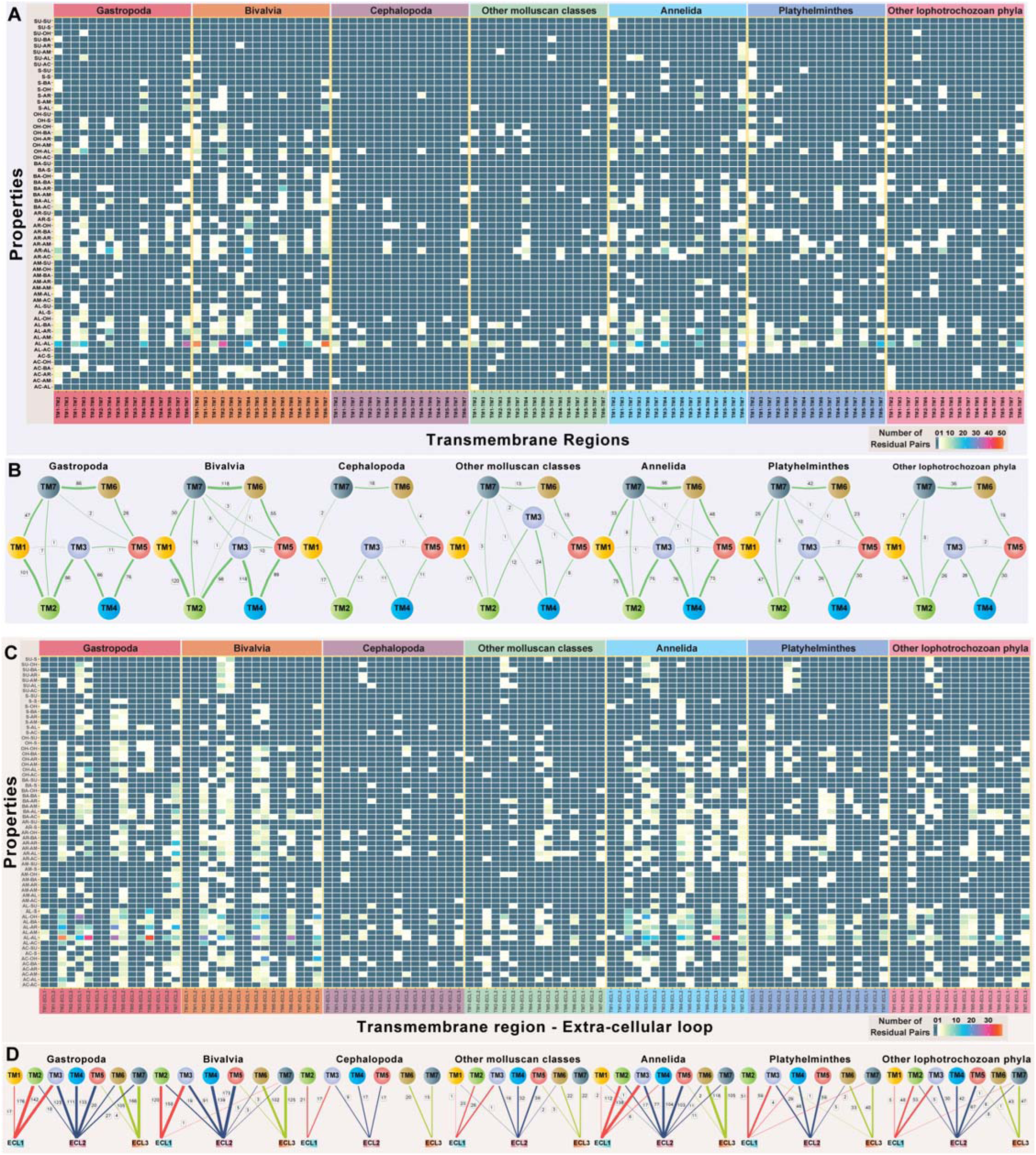
Diversity of residue-contact networks within and across different phyla. **(A)** Inter-TM residue contact network analysis is shown as a heatmap quantifying residue pairs based on amino acid properties (OH: Hydroxyl; AL: Aliphatic; AR: Aromatic; SU: Sulfur-group containing; S: Small-sized; BA: Basic; AC: Acidic; AM: Amide-group containing) across different phyla. The heatmap color scale is on the right, with cells showing zero residue pairs highlighted in teal. **(B)** Schematic representations of inter-TM contacts in different groups, with each TM represented as circles and lines indicating cumulative contacts. The line width is proportional to the number of contacts, and the count is indicated beside each line. **(C)** TM-ECL residue contact network analysis is illustrated as a heatmap quantifying residue pairs categorized by amino acid properties (as in A) across various phyla. The color scheme of the heatmap is the same as panel A. **(D)** Schematic representations of the TM-ECL contact network, with circles for TM and rectangles for ECL. Lines between a circle and a rectangle indicate cumulative contacts across different phyla. The line width corresponds to the number of contacts, with the count indicated beside each line.

Similar to inter-TM contacts, TM-ECL interactions varied significantly in both the number and types of residue pairs across phyla (Fig. 10C and D, table S2). Besides aliphatic-aromatic and aliphatic-aliphatic pairs, TM-ECL interactions predominantly featured basic-basic, basic-hydroxyl, and aliphatic-hydroxyl pairings involving charged or polar residues. Interestingly, no conserved disulfide bridges were found across all phyla in TM-ECL interactions, including the absence of the commonly observed TM3-ECL2 linkage. While some phyla compensated with intra-loop disulfide bridges, gastropods, cephalopods, and other molluscan phyla notably lacked such connections (fig. S2C). Intra-loop disulfide linkages were limited to ECL3 in annelids and ECL2 in bivalves. The absence of disulfide bridges in longer ECL2 suggests significant conformational flexibility, crucial for ligand binding and pocket accessibility, potentially influencing chemosensory receptor interactions.

## DISCUSSION

In this study, we present the first detailed classification of the lineage-specifically expanded subsets of the Class-A GPCR repertoire across 238 lophotrochozoan genomes spanning 8 distinct phyla and provide a comprehensive overview of their evolution, diversification, and correlations with ecological adaptations.

A striking and consistent finding from the comprehensive pan-genomic survey and analysis is that the lineage-specific expansions and reductions align closely with the habitats, ecological niches, and lifestyles of these species. For example, species adapted to extreme environments like deep-sea hydrothermal vents and those with parasitic lifestyles, such as freshwater leeches and parasitic Platyhelminthes, consistently exhibit reduced numbers of Class-A GPCRs and LSEs across various phyla. Similarly, lineages with alternative chemosensory mechanisms, such as cephalopods (*48, 49, 88–90*), and those emphasizing sessile behaviors like certain limpets (*41*), show decreased GPCR and LSE counts. Excluding deep-sea and species with specific lifestyle adaptations, taxa inhabiting various coastal, freshwater, and terrestrial habitats, as well as those with advanced sensory structures across different phyla, typically demonstrate large and divergent LSEs. Examples include shell-less sea slugs, predatory neogastropods, freshwater and terrestrial snails, marine and terrestrial bivalves and annelids, and nemerteans, among others. These observations, together with no significant and consistent role of WGD events in the evolution of these LSEs, illustrate how lifestyle, habitat, and ecological adaptations have shaped the evolutionary dynamics of these expansions and reductions. Further, the diversifications of these LSEs are underscored by the high sequence heterogeneity, signatures of positive selection, and overall conformational flexibility in the ligand-binding pockets, highlighting the adaptations of these receptors to interact with diverse environmental signals.

In summary, our large-scale comparative analysis within this extensive and previously understudied protostome clade reveals a prevalent phenomenon of GPCR-mediated chemoreception akin to its pronounced role in deuterostome lineages. The presented datasets establish a foundational understanding of GPCR-mediated chemoreception in diverse lophotrochozoans, while the methodologies employed offer a valuable resource for future studies, leveraging increased genome accessibility and enhanced assemblies. We propose that our approach is applicable to other major invertebrate lineages, including non-bilaterian metazoans.

## MATERIALS AND METHODS

### Genome Data and the Assessment of the Completeness of the Analyzed Dataset

We sourced all available genomes of lophotrochozoans from various public databases, including the National Center for Biotechnology Information (NCBI) (RRID: SCR_006472), MolluscaDB (http://mgbase.qnlm.ac), Wormbase Parasite (https://parasite.wormbase.org/) (RRID: SCR_003098), MGbase (https://marinegenomics.oist.jp/), CephRes (https://www.cephalopodresearch.org/), and ArgoBase (https://cell-innovation.nig.ac.jp/Aargo/). A total of 261 genomes were downloaded. Of these, 22 genomes were excluded due to poor assemblies, contamination reports, or unreliable results from our preliminary search for Class-A GPCRs, leaving 238 genomes for analysis. The distribution of these genomes is as follows: Gastropoda: 46; Bivalvia: 60; Cephalopoda: 19; other molluscan classes (Solenogastres: 3; Polyplacophora: 4; Monoplacophora: 1; Scaphopoda: 1); Annelida: 32; Platyhelminthes: 59; other lophotrochozoan phyla (Entoprocta: 1; Bryozoa: 5; Phoronida: 3; Brachiopoda: 2; Nemertea: 2). The completeness of these genomes was assessed using BUSCO v5.7.0 (RRID: SCR_015008) with both Metazoa Odb10 and Mollusca Odb10 databases (*91, 92*). The employed BUSCO pipeline used Augustus (RRID: SCR_008417) for gene prediction (*93*), HMMSEARCH as implemented in the HMMER package (RRID: SCR_005305) to detect homologous sequences (*94*), and TBLASTN (RRID: SCR_011822) to align protein with the nucleotide sequences, aiding in the identification of conserved genes (*95*). Of the 238 genomes analyzed, 109 had scores above 90%, with 63 exceeding 95%. Additionally, 33 genomes scored over 80%, and 8 scored between 70-80%. The remaining 88 genomes scored less than 70%; however, 55 of these genomes are from parasitic Platyhelminthes, which exhibit highly reduced genome sizes due to their adaptations towards parasitic lifestyles. Notably, all these genomes are represented as “full” in NCBI, with many being chromosomal-level assemblies (Supplementary Table S1). This suggests that the BUSCO scores are likely biased for these genomes that have undergone significant gene losses and reduction in size. Additionally, the available Odb10 database has a very limited representation of the lophotrochozoan genomes, with only two genomes from Mollusca and one from Annelida, leading to potential clade-specific biases. Such biases were previously documented in genome reports of some of these taxa. Therefore, we retained these genomes for our analysis based on the following criteria: (i) if they are described as full genomes with high coverage in NCBI; (ii) if their Class-A GPCR counts are comparable to those in genomes of the same genus or family with BUSCO completeness scores above 80% or 90% (Supplementary Table S1).

### Sequence Analysis

Sequence similarity searches were conducted using the NCBI BLAST+ (RRID: SCR_004870) and HMMER3 (RRID: SCR_005305) packages (*94, 95*). BLAST searches were performed against the desired databases using BLASTP (RRID: SCR_001010) and PSI-BLAST programs (*95*). Sensitive HMM-based sequence searches were conducted with HMMSERCH and JACKHMMER programs against the proteomes and the compiled database of translated ORFs using the HMMER3 package (*94*). ORFs were predicted and translated in all six reading frames using the NCBI ORFfinder (RRID: SCR_016643) or the getorf program (RRID: SCR_008493) from the EMBOSS package (*96*). HMMBUILD was used to build HMM profiles from manually inspected and corrected seed MSA (Multiple Sequence Alignment), and HMMSCAN was utilized to search sequences against our custom-built HMM database. A plurality-rule consensus sequence was generated from every HMM profile using the HMMemit program by selecting the maximum probability residue at each match state (*94*). HMM-based profile-profile searches were performed using the HHpred program (RRID: SCR_010276) (*97*) against the HMMs derived from PDB (RRID: SCR_012820) and Pfam (RRID: SCR_004726) models (*98*). HHblits against UniRef30 (RRID: SCR_010646) were used as the default method for MSA generation in HHpred, along with other default parameters (*97, 99*). All MSAs analyzed in this study were constructed using the programs Kalign (V2) (RRID: SCR_011810) and MAFFT (RRID: SCR_011811), with manual adjustments based on profile-profile searches against PDB, structural alignments, and predicted 3D structure models (*100, 101*). Removal of redundant sequences and sequence-based clustering was performed using CD-HIT (RRID: SCR_007105) and BLASTCLUST (RRID: SCR_016641) programs (https://www.ncbi.nlm.nih.gov/Web/Newsltr/Spring04/blastlab.html) (*102*).

### Structure Analysis

The JPred (V4) program (RRID: SCR_016504) was utilized to predict secondary structures from the generated multiple sequence alignments (*103*). Each MSA served as a reference model to obtain a 3D structure using AlphaFold2 (AF2) (*104*). The predicted secondary structure boundaries from Jpred4 were compared with the AF2 models for verification. Both the AF2 models and the secondary structure boundaries from Jpred4 were utilized to map the regions forming the ligand-binding groove of the 7TM domain. The figures showing structural visualization and 3D structure rendering were created using the PyMOL Molecular Graphics System, Version 3.0 Schrödinger, LLC (http://www.pymol.org) (RRID: SCR_000305).

### Phylogenetic Analysis

Phylogenetic relationships were inferred using an approximate maximum likelihood (ML) method in the FastTree program (RRID: SCR_015501) (*105*). Local support values were estimated accordingly. To improve topology accuracy, the number of minimum-evolution subtree-prune-regraft (SPR) moves in FastTree was increased to 4 (-spr 4), and the options ’-mlacc’ and ’-slownni’ were used for more exhaustive ML nearest-neighbor interchanges (NNIs). Additionally, phylogenetic tree topologies were derived using ML methods based on the edge-linked partition model in the IQ-TREE2 software (RRID: SCR_017254) (*106*). The best-fit substitution models were determined using the ModelFinder program as implemented in IQ-TREE2 software (*107*). Branch supports were obtained using the ultrafast bootstrap method (1000 replicates) in IQ-TREE2 (*108*). Phylogenetic trees were rendered using the FigTree Software (RRID: SCR_008515) (https://github.com/rambaut/figtree).

### Orthology Inference using OrthoFinder

The OrthoFinder approach (RRID: SCR_017118) that integrates phylogenetic inference of orthologs using gene trees and species trees was utilized to comprehensively identify orthogroups and conserved orthologs across datasets at various taxonomic levels (*109*). The OrthoFinder analysis was conducted using the following settings: BLAST for sequence search, MAFFT for alignment, IQTREE2 for tree inference, STAG for species tree inference (*110*), and MCL clustering with a default inflation parameter of 1.5 (RRID: SCR_024109) (*111*). The results from the OrthoFinder program included an all-inclusive reconstruction of orthogroups, orthologs, gene trees, resolved gene trees, rooted species trees, gene duplication events, and statistics of the overall comparative genomics analysis. These included the number of one-to-one orthologs between each pair of species, the count of orthologs in one-to-many relationships (indicating gene duplication in one of the two lineages post-speciation), and the number of orthologs in many-to-many relationships for each species pair (reflecting gene duplication events in both lineages post-speciation). The inferred datasets and matrices were visualized and plotted using the ggplot2 package (https://ggplot2.tidyverse.org) (RRID: SCR_014601) in R language.

### Curation of Class-A GPCR Repertoires

To assemble the Class-A GPCR repertoire across Lophotrochozoa, we primarily constructed a carefully curated HMM database by sourcing all Class-A GPCR HMM profiles from the Pfam database. Furthermore, we obtained the following: (i) previously curated Class-A GPCR repertoires from various taxa; (ii) clusters of chemosensory GPCRs that have expanded specifically within lineages across diverse metazoan taxa; and (iii) functional olfactory receptors from different vertebrate taxa. Subsequently, separate HMM profiles were constructed using the HMMBUILD program of the HMMER3 package from manually inspected seed alignments of these datasets (*94*). In total, 110 distinct HMM profiles were employed as search seeds using HMMSEARCH and JACKHMMER programs against the proteomes to retrieve all Class-A GPCRs within each analyzed taxon (*94*). Hits obtained from these separate searches, meeting the default inclusion threshold (0.01), were considered putative Class-A GPCRs and were subjected to further scrutiny (see below). Independently, TBLASTN searches were conducted with the retrieved Class-A GPCRs against corresponding genomes to confirm the presence of any additional sequences not represented in the proteome. Likewise, TBLASTN methods were applied to retrieve all Class-A GPCRs for taxa lacking proteome datasets, and translated ORFs were examined for GPCR signatures using the above approach. In parallel, we curated the membrane proteome subset for each proteome using TOPCONS SINGLE and DEEP-TMHMM (RRID: SCR_025039) membrane topology prediction tools (*112, 113*). Protein sequences exhibiting a 7-transmembrane (7TM) topology, with a correction allowance of ± 1TM helical segment to account for any potential erroneous topology predictions, were identified and subsequently validated for the presence of Class-A GPCR signatures. The cross-validation involved conducting PSI-BLAST (*114*) and JACKHAMMER searches (*94*) against the NCBI RefSeq (RRID: SCR_003496) and Swiss-Prot (RRID: SCR_021164) databases and profile-profile HHPRED searches against the HHpred-specific clustered PDB (PDB_mmcif70) and Pfam databases (Pfam-A_v37). Sequences confirmed through prior searches were further scrutinized for typical Class-A GPCR sequence characteristics using taxon-specific multiple sequence alignments. Finally, all non-redundant and non-identical sequences were retained for further analysis. The sequences that failed to meet the criteria were excluded as false positives.

### Identification of LSEs using Integrated Phylogeny and OrthoFinder Approach

We next segregated the Class-A GPCR repertoire in each analyzed taxon into two distinct groups: one comprising the prospective endogenous ligand-binding Class-A GPCRs (such as the amine, peptide, and all other families within Class-A GPCRs) that unequivocally establish one-to-one orthologous relationships across diverse taxa, and the other that had undergone extensive lineage-specific expansions due to local gene duplications—a phenomenon notably observed in chemosensory GPCRs. To achieve this, we adopted a comprehensive hierarchical and recursive phylogenetic tree reconstruction approach. Initially, separate trees were constructed, encompassing numerous taxa at each family level, with a limit of one taxon per genus. This was done to alleviate taxon sampling bias, preventing an overrepresentation of genomes from any specific genus among the 238 genomes representing 168 different genera. Among these, 31 genera had two or more genomes from distinct taxa. To assess the LSE component of the Class-A GPCRs for every member of these 31 genera, the respective family-level trees were replicated, each incorporating one genome at a time from the particular genera. Later, multiple trees were reconstructed at each order and class level by incorporating members from multiple families. The reconstruction of phylogenetic trees at family, order, and class levels was iterated several times using diverse taxonomic combinations to minimize potential biases from taxon sampling during tree reconstruction.

All estimated tree topologies were carefully analyzed, and taxa or lineage-specific paralogous clusters were distinguished from those forming legitimate one-to-one orthologous relationships with sequences from other taxa. The authenticity of these lineage-specific paralogous clusters was verified across all individual trees. Subsequently, from each tree, the clusters forming lineage-specific expansions were separated from subsets forming one-to-one orthologs, and their validation was further ensured using OrthoFinder software to confirm the absence of one-to-one orthologous clusters. For over 95% of the LSEs identified in the phylogenetic trees, OrthoFinder consistently confirmed the absence of one-to-one orthologous relationships. In instances where discrepancies arose and sequences identified as one-to-one orthologs in OrthoFinder analysis but instead grouped within paralogous clusters in the phylogenetic tree, we tested these sequences separately in the phylogenetic trees and deemed them as LSEs if the phylogenetic tree topologies reaffirmed their memberships. Also, orthologs inferred from the OrthoFinder analysis were again cross-verified using the phylogenetic approach as described above, and thus, the LSEs were verified using both phylogeny and OrthoFinder-based approaches (see pipeline in Figure). Moreover, the genomic locations of all LSE-forming clusters were scrutinized to assess their tandem clustering patterns within the studied genome.

### Estimating the Rate of Evolution using Shannon Entropy (H) Analysis

To study the rate of evolution of the categorized LSEs, we estimated the position-wise Shannon entropy (H) for a given multiple-sequence alignment using the following equation:

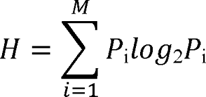

where P is the fraction of residues of the amino acid type, i, and M is the number of amino acid types. The Shannon entropy value for the ith position in the alignment ranges from 0 (only one residue at that position) to 4.32 (all 20 residues equally represented at that position) (*84*). Typically, aligned positions with H > 2 indicate a higher rate of evolution, while H > 2.5 or 3 represent truly fast-evolving positions. Conversely, positions with H < 2 and values closer to 1 are relatively more conserved, while H = 0 indicates completely conserved positions. The entropy values derived thus were analyzed using the R language.

### Structure-Guided Analysis of Ligand-Binding Pockets using Residue Contact Networks

We employed a structure-guided approach to analyze the diversity in ligand-binding pockets of all identified LSEs. First, manually inspected and verified MSAs were generated for all lineage-specifically expanded clusters identified in each analyzed taxon. These MSAs representing the LSEs were then used to define secondary structure boundaries of the 7TM topology using Jpred4. An HMM profile was created using each MSA, and a plurality rule consensus sequence was derived using HMMemit. The obtained consensus sequence served as a representative sequence for generating 3D structures using AF2: a total of 169 structures across all major phyla were generated for further analysis.

Utilizing the MSAs and the AF2-generated 3D structures, we specifically examined the regions located sequentially at the beginning (first ten residues) of TM1, TM3, TM5, and TM7 and at the end (last ten residues) of TM2, TM4, and TM6, positioned structurally at the apex of the 7TM helical bundle, forming the extracellular-facing ligand-binding groove (*115*). We also examined the inner-facing residues within these TM segments that are strategically positioned for ligand interaction. The variation in the length and specific interactions of the dynamic extracellular loops (TM2-TM3:ECL1, TM4-TM5:ECL2, and TM6-TM7:ECL3) with the extracellular-facing TM-segments were also examined, as the flexible loops play a crucial role for determining the pocket shape, overall hydrophobicity, and ligand accessibility (*85, 116, 117*).

For all screened residues from the previous step, residue contact networks for static contacts were estimated, analyzing all residual pairs within intra-TM, inter-TM, intra-ECL, and TM-ECL regions using the GetContacts program (https://getcontacts.github.io/). Before conducting the GetContacts analysis, hydrogen atoms were added to AF2-predicted 3D structures using the Reduce program to improve energy minimization and accurately represent hydrogen bonding and electrostatic interactions, crucial for estimating van der Waals forces and other non-covalent interactions. For each residual pair, the GetContacts program estimates different types of interactions such as the salt bridge, pi-cation bond, pi-stacked bond, t-stacking bond, hydrogen bond, and van der Waals forces and hydrophobic interactions. To summarise, we also categorized each residual pair based on the amino-acid properties involved, such as hydroxyl, aliphatic, aromatic, sulfur-containing, small-sized, basic, acidic, and others. Finally, the number of residual pairs in each category was estimated for intra-TM, inter-TM, intra-ECL, and TM-ECL regions. For example, the number of residual pairs falling in each category, such as “aromatic-hydroxyl (AR-OH)”, Basic-Acidic (BA-AC), and so on within each of these regions were plotted using the ggplot2 package in R. Additionally, inter-TM and TM-ECL contacts were visualized in a network plot using the igraph package in R.

### Genomic Analysis and Clustering Patterns of LSEs

To analyze the genomic context and clustering patterns of LSEs, we obtained the genomic coordinates (in nucleotide base pairs) of all identified LSE sequences. For taxa with both genomic and proteome datasets, we mapped protein accessions to the corresponding GeneID in the NCBI Nucleotide database or the relevant source database to obtain precise coordinates. For taxa with only genomic datasets, predicted gene coordinates were estimated using getorf or ORFfinder and then verified using TBLASTN by mapping the ORFs back to the genome. Using the genomic coordinates, we generated schematic gene scaffolds showing tandem clustering of the LSEs with the ggplot2 package in R. The obtained coordinates are shown as MB base pairs.

### Classification of LSE Members into Full-Length, Truncated, and Pseudogenes

The extracted LSE genes were categorized as follows: Full-Length (FL) if they contained all 7TMs with intact N and C terminals, and truncated if they had 6TMs but lacked portions of the first or last TMs. Sequences with less than 6TMs were excluded from the analysis and not counted among truncated genes. The estimation used automated TM prediction methods, such as TMHMM and DEEP-TMHHM (*113, 118*), for 7TM topology prediction, followed by manual inspection of MSA for each LSE. Pseudogenes were identified using the Pseudogene pipeline script (https://github.com/ShiuLab/PseudogenePipeline) by examining for the presence of premature stop or frameshift mutations, and genes with at least one stop codon and frameshift mutation were categorized as pseudogenes.

## Statistical Analysis

We assessed the normality of the dataset using the Shapiro-Wilk test and by evaluating the Q-Q (quantile-quantile) plot. The homogeneity of variances was determined using Levene’s test. For non-normally distributed or heterogeneous data, we employed the Wilcoxon rank sum test (Mann-Whitney U test) to determine whether there is a statistically significant difference between the distributions of two independent groups and the Kruskal-Wallis rank sum test to determine whether there is a statistically significant difference between three or more independent groups. Kruskal-Wallis rank sum test was followed by Dunn’s post-hoc test with Bonferroni correction for multiple groups. When the data met normality and homogeneity assumptions, we used the two-sample t-test for two independent groups and one-way ANOVA, followed by Tukey’s Honestly Significant Difference (HSD) test for multiple groups. A p-value of less than 0.05 was considered significant. The tests were conducted in R, using the dplyr (RRID: SCR_016708), car (RRID: SCR_022137), and FSA packages and then visualized using ggplot2.

## Author contributions

Conceptualization: AK

Methodology: RN and AK

Investigation: RN, BP, RS, and AK

Visualization: RN, BP, RS, and AK

Supervision: AK

Writing—original draft: AK with contribution from BP and RN

Writing—review and editing: AK with input from all authors

## Data and materials availability

All data needed to evaluate the conclusions in the paper are present in the paper and/or the Supplementary Materials. Fasta sequences, structure models, phylogenetic trees, and raw data files for all the analyses conducted in this study are available in various formats in Zenodo (10.5281/zenodo.12736167).

---

**Fig. S1.**
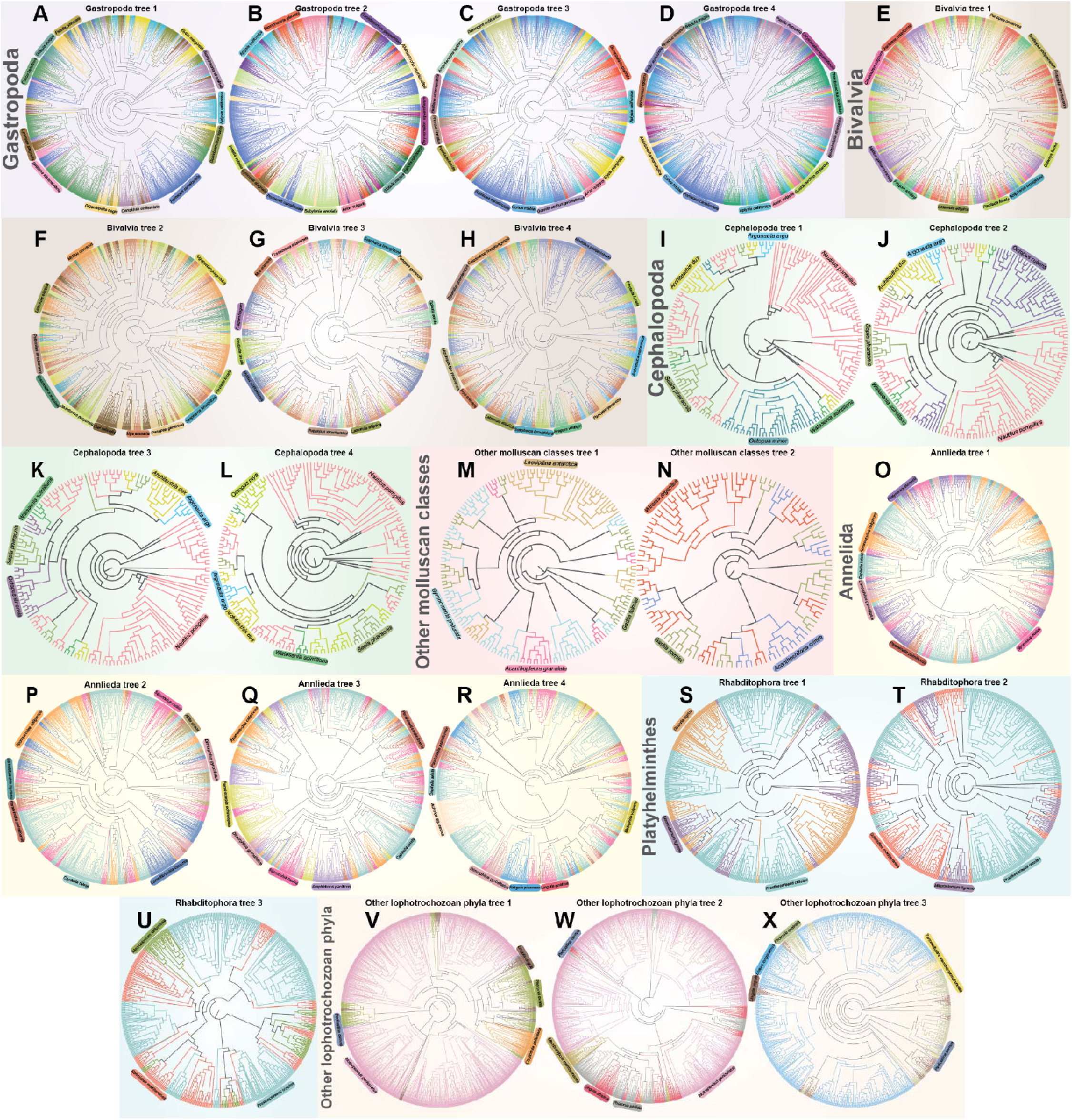
**(A-G)** Maximum-likelihood (ML) tree illustrating lineage-specific clustering of class-A GPCRs from representative taxa across various orders. Multiple iterations of tree construction were performed using different taxa combinations to mitigate bias from over- or under-representation of particular taxa. The taxa-specific phylogenetic trees are divided as follows: **(A)** Gastropoda; **(B)** Bivalvia; **(C)** Cephalopoda; **(D)** other molluscan classes, including Solenogastres, Polyplacophora, Monoplacophora, and Scaphopoda; **(E)** Annelida; **(F)** Rhabditophora (Platyhelminthes); **(G)** other lophotrochozoan phyla, including Entoprocta, Bryozoa, Phoronida, Brachiopoda, and Nemertea. Taxa names are labeled next to their respective clusters.

**Fig. S2.**
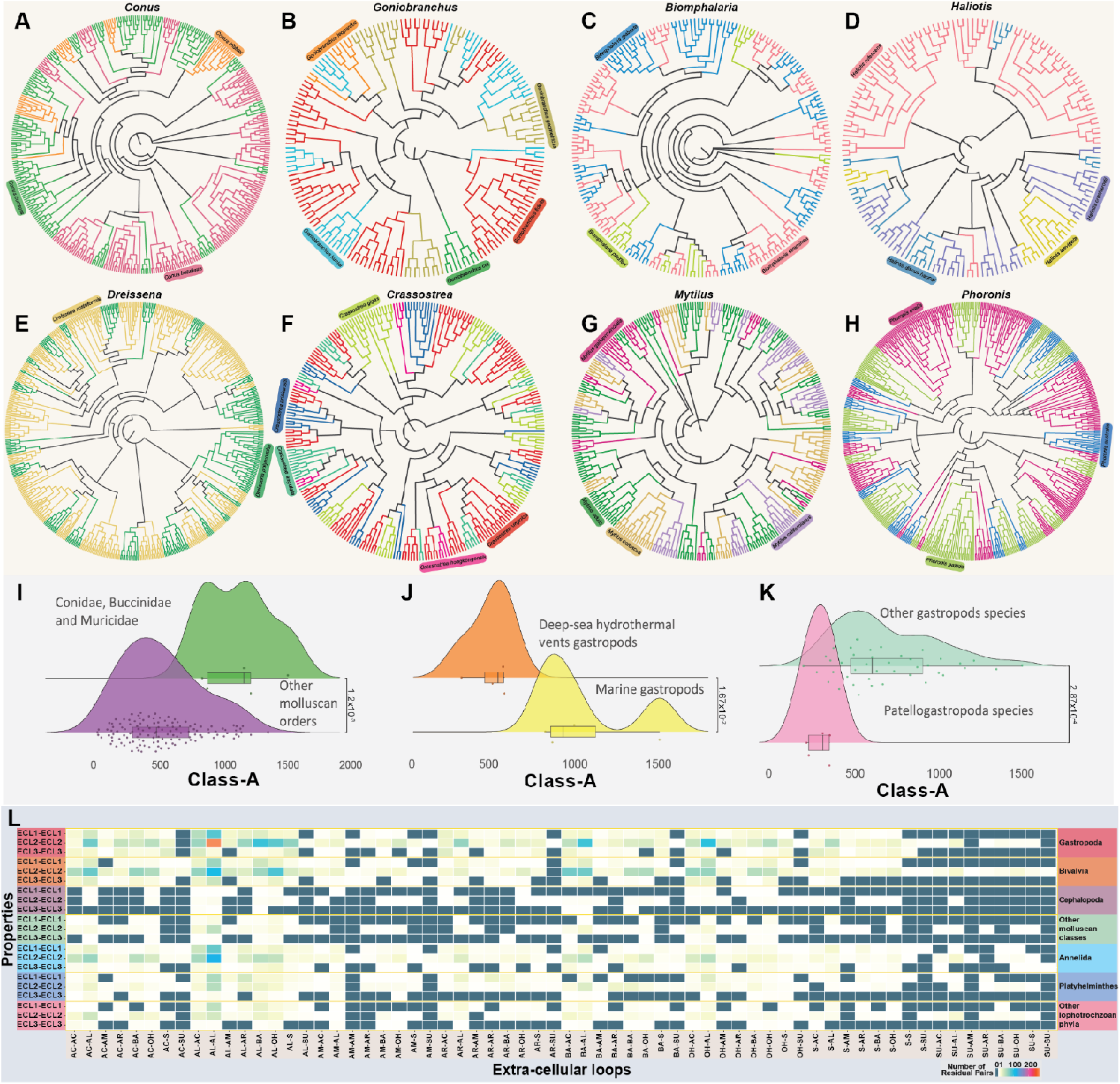
**(A)** Maximum-likelihood (ML) tree illustrating lineage-specific clustering of class-A GPCRs from intra-genus taxa across various orders. The genera are as follows: (i) Goniobranchus; (ii) Conus; (iii) Biomphalaria; (iv) Haliotis; (v) Dreissena; (vi) Crassostrea; (vii) Mytilus; (viii) Phoronis. Taxa names are labeled next to their respective clusters. **(B)** Statistical analysis reveals significant differences in class-A GPCR counts: between the order Neogastropoda (families Buccinidae, Conidae, and Muricidae) and the rest of the molluscan orders (W = 599) (left), between taxa inhabiting deep-sea hydrothermal vents and marine gastropods (t = -3.29; df = 6) (center), and between species in the order Patellogastropoda and other gastropod species (W = 12) (right). The Wilcoxon rank sum test was used for (i) and (iii), and the two-sample t-test for equal variances was used for (ii), with p < 0.05 considered significant. **(C)** Contact network analysis is depicted as a heatmap quantifying the residual pairs between intra-ECL categorized based on their amino acid properties (OH: Hydroxyl; AL: Aliphatic; AR: Aromatic; SU: Sulfur-containing; S: Small-sized; BA: Basic; AC: Acidic; AM: Amide-group containing) across different phyla. The heatmap color scale is on the right, with cells showing zero residue pairs highlighted in teal.

